# Exploring structure-function relationships in engineered receptor performance using computational structure prediction

**DOI:** 10.1101/2024.11.07.622438

**Authors:** William K. Corcoran, Amparo Cosio, Hailey I. Edelstein, Joshua N. Leonard

## Abstract

Engineered receptors play increasingly important roles in transformative cell-based therapies. However, the structural mechanisms that drive differences in performance across receptor designs are often poorly understood. Recent advances in protein structural prediction tools have enabled the modeling of virtually any user-defined protein, but how these tools might build understanding of engineered receptors has yet to be fully explored. In this study, we employed structural modeling tools to perform post hoc analyses to investigate whether predicted structural features might explain observed functional variation. We selected a recently reported library of receptors derived from natural cytokine receptors as a case study, generated structural models, and from these predictions quantified a set of structural features that plausibly impact receptor performance. Encouragingly, for a subset of receptors, structural features explained considerable variation in performance, and trends were largely conserved across structurally diverse receptor sets. This work indicates potential for structure prediction-guided synthetic receptor engineering.

## INTRODUCTION

Engineered cell therapies are a burgeoning approach for treating diseases ranging from cancer^1,2^ to autoimmune diseases^3,4^. A key benefit of engineered cell therapies is their ability to elicit a targeted therapeutic response at the site of disease. This precision is commonly conferred by engineering cells to express synthetic receptors, which transduce the detection of user-defined cues into the induction of therapeutic responses. A diverse repertoire of engineered receptors now exists^5^, including chimeric antigen receptors (CARs)^6^, synthetic notch receptors (synNotch)^7,8^, synthetic intramembrane proteolysis receptors (SNIPRs)^9^, generalized extracellular molecules sensors (GEMS)^10^, and modular extracellular sensor architecture receptors (MESA)^11,12^. Each platform confers unique functions, and all employ distinct mechanisms, but a common trait is that the structure of the receptor is generally poorly understood and impacts performance in ways that are often subtle, substantial, and difficult to predict, necessitating iterative, empirical refinement.^13–15^ Generating tools and workflows for improving our understanding of the relationships between engineered receptor structure and function could streamline the process of generating new, high-performing synthetic receptors for diverse applications.

Elucidating the structure of natural and engineered receptors poses unique challenges, but emerging tools may help to address these needs. Many engineered receptors are single-pass transmembrane proteins, which are generally a particularly challenging class of proteins for structural characterization as they are often relatively large, contain a hydrophobic transmembrane domain connecting their extracellular and intracellular components, and can adopt multiple conformations.^16,17^ Given this complexity and the time and effort required to resolve protein structures, we have some structural information about synthetic receptor subdomains^18^ but lack complete structures for the vast majority of engineered receptors. A promising recent advance—and subject of the 2024 Nobel Prize in Chemistry to Baker, Hassabis, and Jumper^19^—is the emergence of protein structural prediction tools such as AlphaFold^20,21^, AlphaFold-Multimer^22^, and RoseTTAFold^23^, which enable structural predictions of proteins for which there is no experimentally observed structure. Recently, these tools have been deployed to construct complete models of natural single-pass transmembrane receptors^24^ and this approach has been incorporated into the engineering process for modifying receptors^25^. Whether these tools can help to build understanding of structure-function relationships for engineered receptors represents an exciting and largely unexplored frontier.

In this study, we investigated the use of computational protein structure prediction tools to generate new insights into a pre-existing dataset comprising empirical characterizations of a library of synthetic receptors. We employed a recent investigation focused on conversion of natural human cytokine receptor ectodomains into engineered MESA receptors (termed NatE MESA).^26^ MESA is a two-chain system in which extracellular ligand binding-induced heterodimerization of these chains drives intracellular reconstitution of a split tobacco etch virus protease (TEVp), which releases a sequestered transcription factor to drive expression of an output transgene (**Fig. 1A**).^27^ This dataset is of particular interest for the current study because this ensemble of receptors encompasses variation in both structure and performance, suggesting that we might be able to discern which, if any, structural variables correlate with performance metrics of interest. In addition, this family of receptors is based upon domains derived from natural receptors, for which we hypothesized that structural prediction might be most feasible given the reliance of existing tools on sequence alignment with known proteins. Nonetheless, the current protein databank (PDB) lacks complete structures for many combinations of receptor domains and ligands explored in this study. Therefore, computational structural prediction could be uniquely valuable for explaining observed variations in a way that exceeds both intuition and existing structural knowledge. Encouragingly, we found that a considerable amount of variation in NatE MESA performance could be explained by variation in the structure of these receptors (**Fig. 1B**). We were able to attribute this variation to several key independent variables (features), and many relationships between structural variables and performance were consistent across NatE MESA receptor families. This work provides a general workflow that may help identify structure-function relationships for both natural and engineered receptors to increase our understanding of receptor biology and guide future engineering.

**Figure 1.**
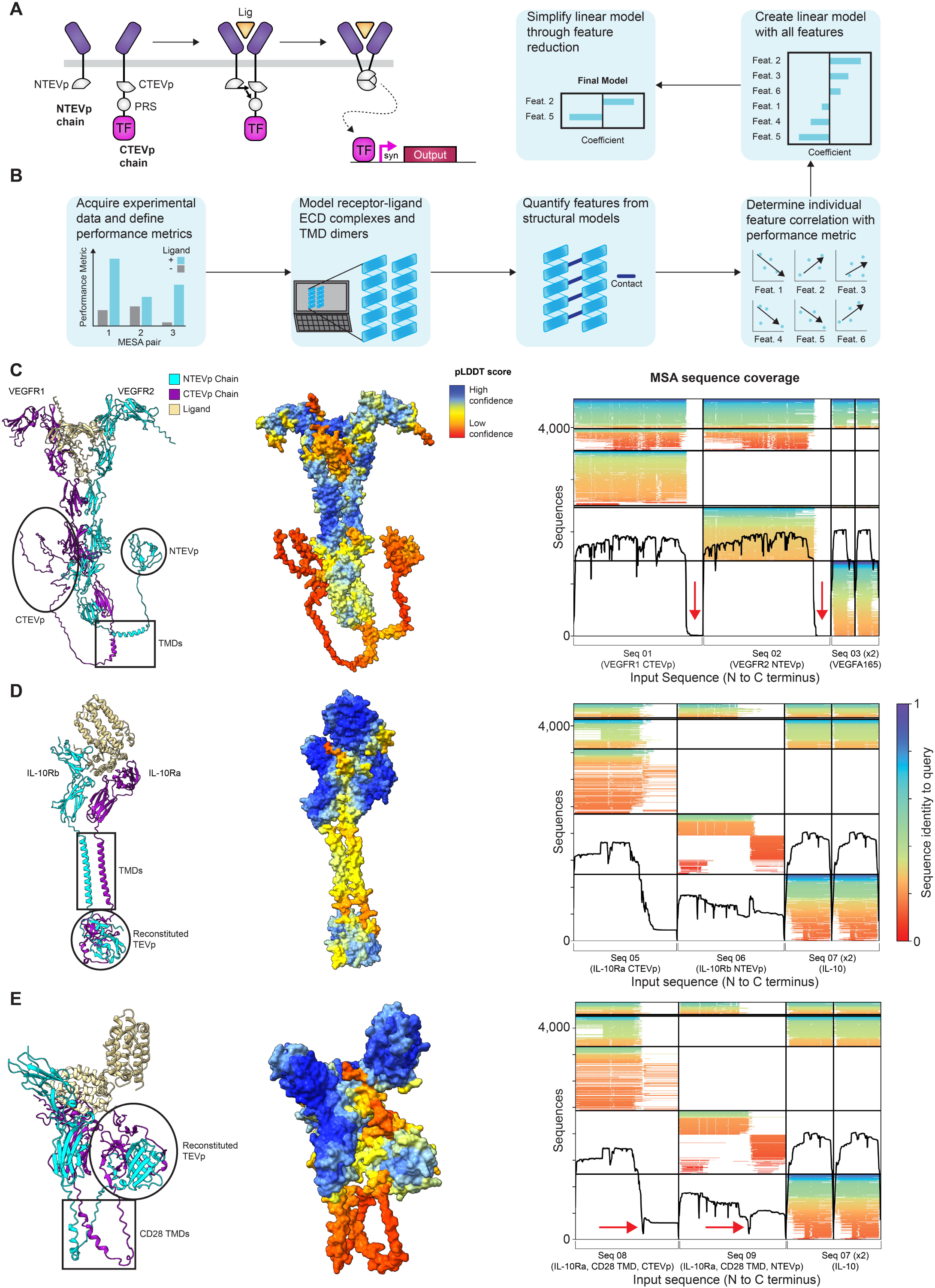
Establishing methods for post hoc analysis of MESA structure-function relationships. (**A**) Schematic representation of MESA receptor signaling mechanism. Ligand binding induces dimerization of the NTEVp and CTEVp chains, leading to reconstitution of TEVp and autoproteolytic release of a sequestered transcription factor to drive output gene expression. (**B**) Overview of the workflow used in this investigation. (**C**) ColabFold-generated model of the complete VEGFR2 NTEVp and VEGFR1 CTEVp chains in complex with a VEGFA165 dimer (**Supplementary Data 1, Structure 1**). The NTEVp and CTEVp domains do not appear reconstituted in this structure (left), receive low pLDDT scores (middle), and lack sequence coverage in the MSA in these regions (right, red arrows). Vertical lines in the sequence coverage plot delineate individual protein sequences included in the query (i.e., receptor chains or ligands in the complex), and horizontal lines delineate distinct ColabFold multiple sequence alignment outputs, where each alignment includes one or more query protein sequences. (**D**) ColabFold-generated model of the complete IL-10Rb NTEVp and IL-10Ra CTEVp chains in complex with an IL-10 dimer (**Supplementary Data 1, Structure 2**). The reconstituted TEVp (left) has a higher pLDDT score (middle) compared to the VEGFR complex in **B**, and the sequence coverage plot (right) shows coverage across the NTEVp and CTEVp regions of their respective chains. (**E**) ColabFold-generated model of the complete IL-10Rb NTEVp and IL10Ra CTEVp chains, each of which includes a CD28 TMD in place of the native TMD (depicted in **D**) (**Supplementary Data 1, Structure 3**). While TEVp appears reconstituted (left), the TMDs appear curved, with low pLDDT scores (middle), and TEVp is folded upward near the ECDs, which is not physically realistic. Sequence coverage in the MSA drops at the TMDs (right, red arrows). Abbreviations: N-terminal half of Tobacco Etch Virus Protease (NTEVp), C-terminal half of Tobacco Etch Virus Protease (CTEVp), protease recognition sequence (PRS), transcription factor (TF), synthetic promoter (syn), ligand (lig), transmembrane domain (TMD), feature (Feat.), vascular endothelial growth factor receptor 1 (VEGFR1), vascular endothelial growth factor receptor 2 (VEGFR2), vascular endothelial growth factor (VEGF), interleukin 10 receptor a (IL-10Ra), interleukin 10 receptor b (IL-10Rb), interleukin 10 (IL-10).

## RESULTS

### Generation of MESA structural models using ColabFold and PREDDIMER

As a first step toward investigating structure-function relationships for MESA, we employed ColabFold^28^ to generate structural predictions of reported NatE MESA variants.^28^ ColabFold is an accessible structure prediction tool providing efficient multiple sequence alignment (MSA) search and a user-friendly online notebook to generate structural predictions using AlphaFold2^20^ and AlphaFold- Multimer^22^. We generated models of the four distinct receptor families investigated in the NatE MESA study, named based upon their ligand: vascular endothelial growth factor (VEGF MESA), tumor necrosis factor (TNF MESA), interleukin 10 (IL-10 MESA), and transforming growth factor beta (TGFβ MESA). ColabFold generates up to five unique structural models of a protein complex (or protein) from the user-defined amino acid sequence and ranks models of protein complexes based upon their predicted template modeling (pTM) and interface-predicted template modeling (ipTM) scores.^29^ Both confidence scores range between 0 and 1, with higher values indicating greater confidence in the structure of the protein complex. Using the highest-ranked model, we then assessed individual amino acids and specific domains based upon the predicted local distance difference (pLDDT) score, which captures the confidence in predicted local structures, and scores range from 0 to 100 (scores above 90 are high confidence, and scores below 50 are low confidence). Given that NatE MESA receptors comprise natural domains combined in novel ways with synthetic elements, we first examined how these compositional aspects impact the confidence of structural predictions.

We first attempted an ambitious approach, modeling several entire receptor-ligand complexes. Each receptor chain comprises an ectodomain (ECD), transmembrane domain (TMD), and intracellular domain (ICD), which includes the respective TEVp component for each chain—the N-terminal half (NTEVp) or the C-terminal half (CTEVp).Individual receptor chains in the NatE study were designed by pairing each natural receptor ECD with either the natural TMD or a murine CD28 TMD and either CTEVp or NTEVp as the ICD. We first predicted the structure of VEGF MESA, which includes two chains (one NTEVp and one CTEVp) bound to a homodimeric VEGF ligand (**Fig. 1C**). These models yielded relatively high pLDDT scores through the ECD, but the score dropped for the TEVp components, which appeared largely unstructured. Analysis of the sequence coverage plot produced by ColabFold revealed no sequences aligning to the NTEVp or CTEVp domains (**Fig. 1C**). Since MSAs are a critical informational input for structure prediction in ColabFold^20,28^, it is possible that inadequate MSA coverage could produce low confidence regions in the predicted structure. We reasoned that the lack of MSA coverage across the TEVp components could be attributed to the large size of the VEGFR ECDs relative to the TEVp components, as the ECDs comprise approximately 80-90% of the total receptor chain. This could cause sequences that align to the TEVp components to receive low alignment scores during the sequence search and to be dropped from the MSA; in this interpretation, the barrier is not a fundamental limitation for sequence alignment but rather a mismatch between how sequence alignment is deployed in ColabFold and the particular chimeric protein structure prediction attempted here. In support of that interpretation, predicting the structure of smaller receptor chains, such as IL-10 MESA, resulted in a completely structured receptor-ligand complex, with reasonably high confidence in both the ECD and TEVp components, and moderate confidence through the TMD (**Fig. 1D**). To further explore the role that chimerism and sequence alignment may play in structure prediction, we next analyzed a similar IL-10 MESA design in which the native TMD was replaced with the CD28 TMD (murine). These new designs yielded low-confidence TMDs along with overall structures that curve upwards and lack plausibility and utility (**Fig. 1E**). We explored improving the quality of the prediction for this complex by constructing a custom MSA with additional sequences covering the CD28 TMD, which were filtered to ensure adequate sequence diversity. However, this change resulted in little improvement in the quality of the structural prediction, which still exhibited curved TMDs (data not shown). Since including the physical constraints imposed by the membrane (in real life) is not possible using current modeling tools, these modeling successes and failures provide some guidance as to how current tools may be best deployed in this investigation.

To enable cross-comparison over all receptors in the four NatE MESA families, we elected to use a generally applicable process in which we modeled ECDs and TMDs as isolated components. Using ColabFold to generate models of receptor ECDs in complex with their ligand resulted in reasonably high confidence metrics for all four receptor families (**Supplementary Table 1**). To model TMD interactions, we selected a tool called PREDDIMER, which is designed specifically to capture TMD biophysics and can accommodate heterotypic pairings.^30,31^ Together, these tools enabled us to generate an ensemble of structural models covering all ECD-ligand complexes and all TMD pairings, and we next investigated potential relationships between predicted structural features and empirical functional performance.

### Separate ECD and TMD models enable the quantification of structural features

To guide our exploration of potential structure-function relationships, we defined a set of candidate structural features that we hypothesized could impact function, and which vary across the receptors modeled. In this framing, features are independent variables and performance metrics are the dependent variables. We first defined two ECD structural features: (i) the distance between the ECD C-termini (ECD distance), as this could affect the ability of the TMDs to dimerize and TEVp to reconstitute (**Fig. 2A**); and (ii) the number of contacts between the two ECD domains (ECD contacts), as variation in the strength of inter-chain interactions could impact the duration and strength of receptor dimerization (**Fig. 2B**). We then defined four TMD features: (i) distance between the C-termini of the TMD models (TMD distance), as this distance could impact TEVp reconstitution (**Fig. 2C**); (ii) the number of contacts between the TMDs (TMD contacts), as we reasoned that the strength of the TMD interaction could impact the duration of dimerization and TEVp reconstitution (**Fig. 2D**); (iii) the angle created between the two dimerized TMD helices (TMD crossing angle), as variation in the TMD crossing angle could impact both the proximity and orientation in which NTEVp and CTEVp interact (**Fig. 2E**); and (iv) TMD exit angle (**Fig. 2F**); as we were unable to generate structural models with receptors containing NTEVp or CTEVp domains, we defined TMD exit angle (the angle measured between vectors oriented along each final residue of a TMD from a PREDDIMER prediction) as a metric that could capture compatibility between TMD orientation and the requirements for reconstituting halves of split TEVp. These metrics form a basis set comprising quantitative (extensive) structural features, which we use alongside categorial features (e.g., the identity of specific domains), as discussed below, as potential independent variables that may contribute to performance.

**Figure 2:**
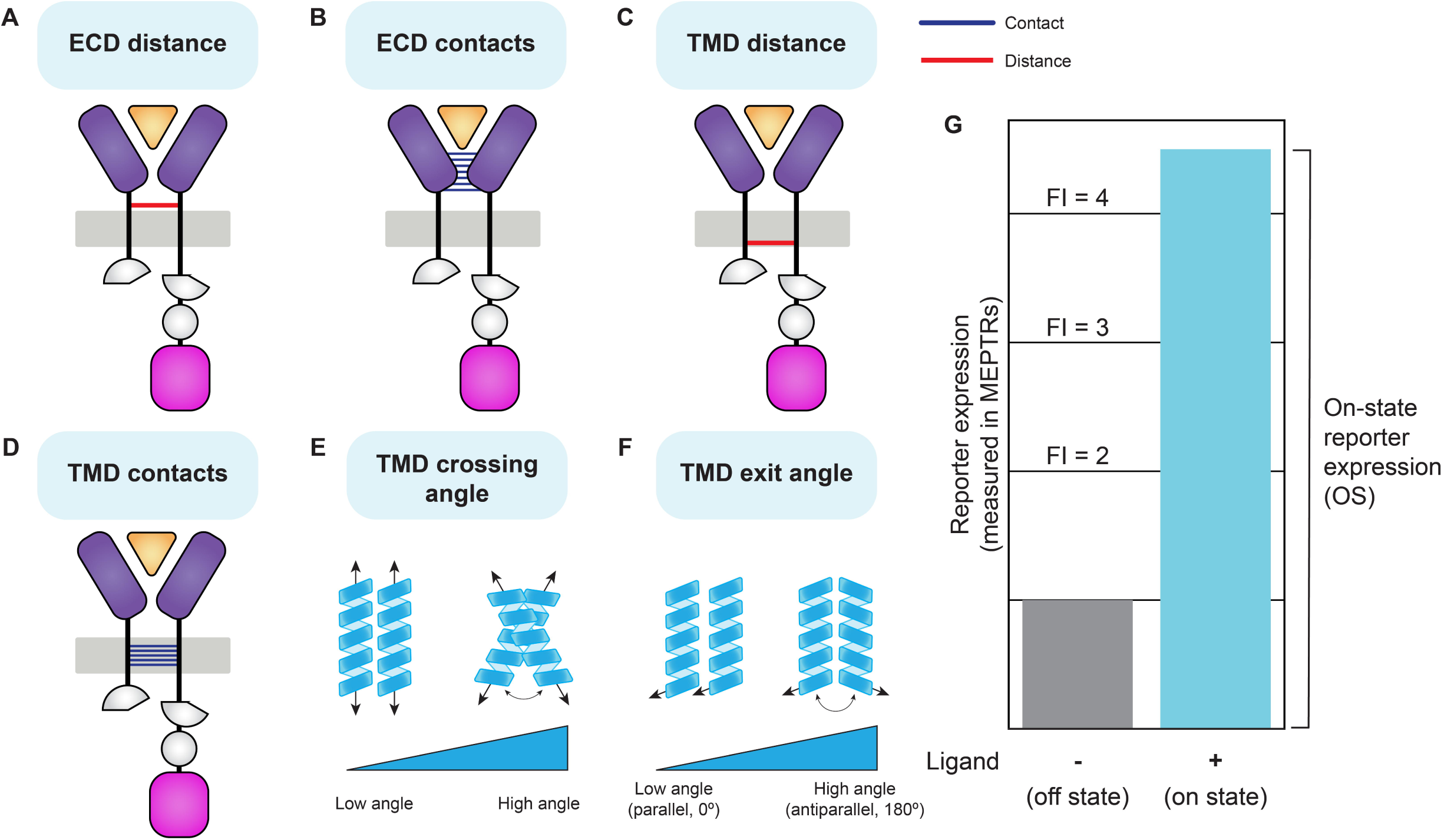
Definition of structural features and performance metrics. (**A-G**) Schematic explaining the structural features (independent variables) and performance metrics (dependent variables) used in structure-function analyses. (**A**) ECD distance: the distance in Å between the Cα of the last residue in each chain predicted to be a part of a secondary structural element (β sheet or α helix). (**B**) ECD contacts: the number of inter-chain contacts between two MESA chains in complex with their cognate ligand. (**C**) TMD distance: the distance in Å between the Cα of the last residue in each TMD. (**D**) TMD contacts: the number of inter-chain contacts between two TMDs. (**E**) TMD crossing angle: calculated using PREDDIMER and defined as the angle created between two TMDs when dimerized. (**F**) TMD exit angle: calculated using TMD models and defined as the angle created between the vectors aligned with the final backbone residues of each TMD. This angle will always be between 0° and 180°. (**G**) Hypothetical data explaining performance metrics: on-state reporter expression (OS) and fold induction (FI). OS is the magnitude of reporter expression when the ligand is present, and FI is the relative magnitude of reporter expression between the ligand-present and ligand-absent conditions. Reporter expression values from the NatE MESA investigation were provided in a standardized unit, molecules of equivalent PE-Texas Red (MEPTRs). Abbreviations: ectodomain (ECD), transmembrane domain (TMD), fold induction (FI), molecules of equivalent PE-Texas Red (MEPTRs).

We next defined quantitative MESA performance metrics to serve as the dependent variables for our structure-function analyses. The two performance metrics most commonly used to evaluate and compare receptor variants, when the output is transcription of a reporter gene, are the on-state reporter expression (OS) and fold induction (FI). OS quantifies the magnitude of the reporter expression when the ligand is present, and FI quantifies the relative reporter expression when the ligand is present versus when the ligand is absent (**Fig. 2G**). For both metrics, higher values indicate better performance. Within the empirical NatE MESA data, variation in OS was greater than variation in FI (**Supplementary Fig. 1A, C**), and thus we focused primarily on OS as a dependent variable that may be likely to capture any structure-function relationships.

### Structural and categorical features explain variation in VEGF MESA performance

To test the hypothesis that differences in predicted protein structure could explain variation in MESA performance, we selected VEGF as the initial case study given the large observed variation in both OS and FI (**Fig. 3A, Supplementary Fig. 2A**). We first manually examined the library of predicted structures. This dataset contains 16 unique receptor pairings and two homodimeric ligands, VEGFA121 and VEGFA165, totaling 32 unique combinations (**Source Data**). Mechanistically, natural VEGFRs can form homo- or heterodimers and primarily signal in their ligand-bound state.^32,33^ We have previously hypothesized that VEGF MESA signals through similar ligand-mediated dimerization, though ligand-independent signaling was evident in some VEGF MESA variants.^26^ Within our structural predictions we observed minimal differences in ECD features as a function of ligand choice (VEGFA121 or VEGFA165), indicating that performance variation observed across these ligands was unlikely to be explained by predicted differences in receptor-ligand complex structure (**Supplementary Fig. 3A, B, D**). However, substantial differences in ECD distance were predicted across the three VEGFR ECD pairings (VEGFR1- VEGFR1, VEGFR1-VEGFR2, VEGFR2-VEGFR2) (**Supplementary Fig. 4A-C**). The VEGFR1-VEGFR2 model has the closest ECD distance at only 26.5Å apart, while the VEGFR2-VEGFR2 ECD model shows a strong divergence between domain 7 (the C-terminal domain) in each chain, resulting in >70Å distance between the C-termini. Of note, this is a distinctly different configuration than was observed in a previously reported VEGFR2 domain 7 dimer (PDB ID: 3KVQ).^34^ It is possible that the structure of the entire ECD dimer differs from that of isolated domain 7 dimers, or perhaps interaction between the domain 7 pair is relatively weak such that these domains were not predicted to interact by ColabFold. We also generated models of the three VEGFR ECD pairings in the ligand-free configuration to assess whether quantifying differences between the predicted ligand-bound and ligand-free (apo) structures could plausibly explain differences in reporter signal between the ligand-free (background) and ligand-added functional evaluations. However, the predicted apo and ligand-bound structures were nearly identical (**Supplementary Fig. 3E**). This suggests a limitation as to how these tools can be used and renders it unlikely that minute differences in predicted ECD structure +/- ligand could explain the sizeable differences in observed reporter expression. Thus, our analysis focused on exploring differences in structural features across the various ligand-bound receptor complexes.

**Figure 3:**
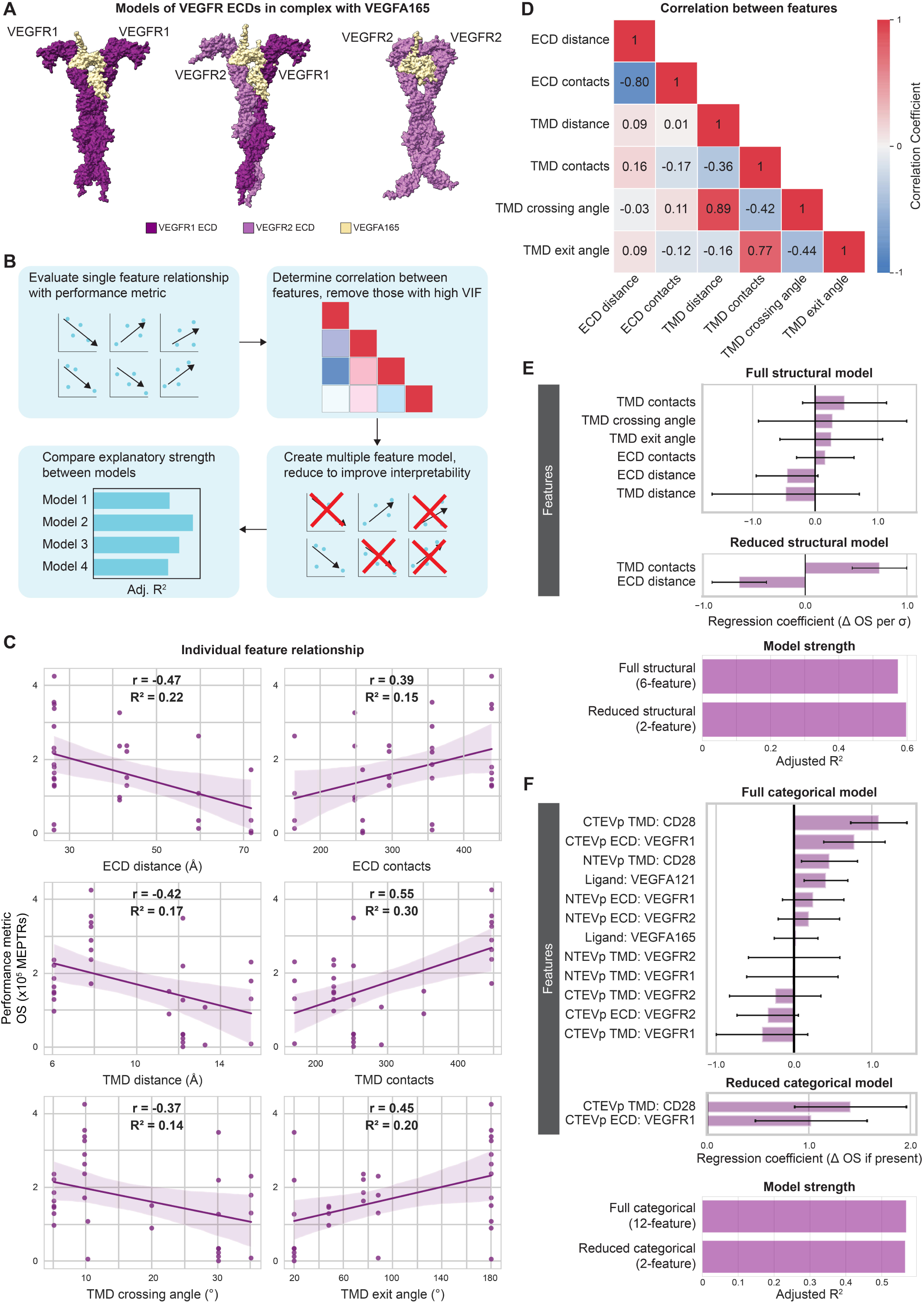
Structural variables explain considerable performance variation in VEGF MESA. (**A**) ColabFold-generated models of VEGFR ECDs in complex with VEGFA165 (tan): VEGFR1-VEGFR1 (left), VEGFR1-VEGFR2 (middle), VEGFR2-VEGFR2 (right) (**Supplementary Data 1, Structures 5, 7, 9**). (**B**) Generalized workflow for identifying key structural features that account for variation in MESA performance. (**C**) Individual structural feature contributions to OS. (**D**) Evaluation of feature independence (Pearson’s correlation coefficients). (**E**) Magnitude of the coefficients for the full (six-feature) and reduced (two-feature) structural models (top), and comparison of the two models’ strengths in explaining variation in OS, as determined by adjusted R^2^ (bottom). Structural feature data were normalized to mean = 0 and standard deviation = 1 prior to analyses. Error bars represent +/- 95% confidence intervals calculated using Student’s t-distribution. (**F**) Magnitude of the coefficients for the full (twelve-feature) and reduced (two-feature) categorical models (top), and comparison of the two models’ strengths in explaining variation in OS, as determined by adjusted R^2^ (bottom). Error bars represent +/- 95% confidence intervals calculated using Student’s t-distribution. Abbreviations: vascular endothelial growth factor receptor (VEGFR), vascular endothelial growth factor receptor 1 (VEGFR1), vascular endothelial growth factor receptor 2 (VEGFR2), vascular endothelial growth factor (VEGF), adjusted R^2^ (Adj. R^2^), Pearson’s correlation coefficient (r), coefficient of determination (R^2^), on-state reporter signal (OS), molecules of equivalent PE-TexasRed (MEPTRs), ectodomain (ECD), transmembrane domain (TMD), cluster of differentiation 28 (CD28), N-terminal half of Tobacco Etch Virus Protease (NTEVp), C-terminal half of Tobacco Etch Virus Protease (CTEVp), change in (Δ), standard deviation (σ).

We next evaluated the strength of the relationship between each structural feature and OS using simple linear regression. We opted to start with simple linear regression over more complicated approaches to feature selection (e.g., random forest or lasso regression) to manage risk of overfitting and build interpretability. Our general approach was to first evaluate the contribution of individual features, then identify redundancies through feature selection, then consider combined contributions of features in complete and reduced statistical models (**Fig. 3B**). Across the entire VEGF dataset, all features exhibited weak correlations with OS and could explain no more than 30% of the variation in OS as determined by the coefficient of determination (R^2^) (**Fig. 3C**). Thus, no individual feature is, alone, a strong predictor of OS. The two distance-based features—ECD distance and TMD distance—exhibited a negative relationship with OS; greater inter-chain distances are associated with lower OS (**Fig. 3C**). This finding is promising, in that this trend concords with prior empirical analyses of distinct MESA structural variants.^35^ Both contact-based features—ECD contacts and TMD contacts—exhibited positive relationships with OS; greater numbers of inter-chain contacts correlate with higher OS (**Fig. 3C**). Larger TMD crossing angles were associated with lower OS, and TMD exit angle analysis indicates that as the final residues of the TMDs faced more opposite directions (closer to 180°), OS tended to increase (**Fig. 3C**).

Next, we evaluated the strength of the relationship between each structural feature and FI. Since the FI data were right-skewed, we first log-transformed the (dependent variable) data to reduce the influence of outliers on the regression models (**Supplementary Fig. 2A**). Similar to relationships with OS, we found that five of the six features explained low to moderate variation in log-transformed FI, while TMD contacts explained nearly half of the variation in log-transformed FI (R^2^ = 0.48) (**Supplementary Fig. 2B**). Both ECD features correlate with the same sign (positive or negative) with log-transformed FI as they did with OS (**Supplementary Fig. 2C**). This combined finding further suggests that shorter ECD distances and greater numbers of ECD contacts may benefit overall VEGF MESA performance. Notably, all TMD features correlate with log-transformed FI with the opposite sign as OS (**Supplementary Fig. 2C**). One plausible explanation for this comparative finding is that TMD features associated with higher OS may also be associated with higher background (off-state) signaling, which in turn reduces FI. Even with the caveat that such relationships are weak, these initial findings are physically plausible, suggest testable hypotheses (i.e., guiding potential future experimental work), and motivated us to continue investigating potential structure-function relationships in these data.

Given that no individual structural feature explained greater than 30% of the variation in VEGF MESA OS, we next evaluated the extent to which combining all features into a single, linear model improved explanatory power. We first scaled all variables to mean = 0, standard variation = 1 to avoid potential influence due to unit or measurement differences. We next explored the potential for collinearity between variables by calculating a correlation matrix (**Fig. 3D**). ECD distance and ECD contacts had a strong relationship (Pearson’s r = -0.8), and multiple TMD features had strong relationships with one another. The strength of these relationships is not necessarily surprising, as for example, an increase in the TMD crossing angle would result in an increase distance between the TMD C-termini (TMD distance). The correlation matrix also revealed weak relationships between ECD variables and TMD variables, indicating that ECD features could explain different sources of variation in MESA performance than do TMD features.

We next explored the explanatory power of the multi-feature model. The full structural model (including all six structural features) had considerably greater explanatory strength as measured by adjusted (Adj.) R^2^ (0.57) than did any of the six individual-feature models (**Fig. 3E**). When considering the explanatory strength of such models, we employed adjusted R^2^ because adding features always improves a model, or at least does not reduce the explanatory strength. If a feature offers very little (or no) improvement, the coefficient for that feature would be at or near zero. Using the adjusted R^2^ metric slightly penalizes the use of additional features, so that one can compare model strength in a manner that depends less on the number of features included. In our analysis, not all features significantly improved the model as determined by each coefficient’s p-value (α = 0.05). Thus, we next simplified the linear model to improve its interpretability, keeping only the essential features that explained OS variation. First, we removed collinear features by determining the variance inflation factor (VIF) for each feature in the group and removed the feature with the highest VIF (threshold: VIF ≥ 10) (**Supplementary Fig. 5A**). We subsequently performed backward elimination of features based on their coefficient’s p-value, removing the feature with the highest p-value one-by-one, arriving at a final model including only features with a statistically significant (α = 0.05) coefficient (**Supplementary Fig. 5B**). This reduced structural model contained only two features, ECD distance and TMD contacts, which together explain 60% of the variation in OS (Adj. R^2^ = 0.60). This reduced model has greater explanatory strength while also providing clarity as to the critical features in explaining variation in VEGF MESA OS.

Given the promising results obtained relating predicted structural features to performance, we next extended this analysis to consider, alternatively, features that capture the identity of specific domain choices used to compose receptor chains. We deemed these “categorical” variables, with the expectation that they might similarly probe information contained in the dataset but with perhaps less generalizability for generating mechanistic insights. Categorical variables may capture properties that are unique to the individual chains included in the MESA receptor pair (e.g., protein expression, membrane localization, protein stability, etc.), which are not directly captured by our structural features. As an example, VEGF MESA pairings with CD28 TMDs tend to produce the highest OS. It is possible that this could result from the high number of contacts between CD28 TMDs, which would be captured by structural feature analysis, but such trends could also result from other factors, such as higher protein expression or clustering driven by CD28 TMDs^26,36^, which are variables not captured by our structural feature model. Further, our structural model does not distinguish differences in chain identity, as determined by the ICDs (NTEVp or CTEVp, which were not included in these structural predictions), although chain identity does impact functional importance. For example, pairing VEGFR2 NTEVp with VEGFR1 CTEVp yielded good experimental performance, while the inverse pair, VEGFR1 NTEVp paired with VEGFR2 CTEVp, yielded worse performance.^26^ We speculate that such differences could reflect changes in the NTEVp-CTEVp orientation upon receptor-ligand complex formation. The structural model does not differentiate between these two pairings, as their structural feature measurements are derived from the same ECD and TMD in a predicted structure. Thus, we expanded our analysis to capture these potentially important aspects of receptor design.

Our categorical model probed the contribution of individual receptor chain components to overall performance. Categorical features were implemented using one-hot encoding, wherein a “1” represents the presence of a component and “0” represents the absence of a particular component on a particular chain. This encoding accounts for whether a component is on the NTEVp or CTEVp chain for each VEGF pairing. The full categorial model (including all 12 one-hot encoded features) explained slightly less variation in OS (Adj. R^2^ = 0.58) than did the two-feature structural model (Adj. R^2^ = 0.60) (**Fig. 3E, F**). We next used backward elimination to retain only statistically significant features, and the resulting reduced categorial model explained a similar amount of variation in OS (Adj. R^2^ =0.57) (**Fig. 3F**). This reduced categorical model comprised two features, each with positive coefficients, indicating that OS is greater for receptors in which the CTEVp chain contains a CD28 TMD and the CTEVp ECD is VEGFR1. Altogether, the reduced categorical model may help to identify domain choices that most impacted performance, but interpretability is limited, and such correlations do not readily identify generalizable trends that may apply across NatE MESA receptor families.

We next explored whether the reduced (two-feature) structural model and the reduced (two-feature) categorical model captured similar information by creating a combined model. We considered the possibility for informational overlap between the models as MESA pairings with at least one VEGFR1 ECD tended to have closer ECD distance and CD28 TMD pairings tended to have a higher number of contacts. The combined linear model explained 79% of the variation within OS (Adj. R^2^ =0.79), and all four features were statistically significant (**Supplementary Fig. 6A-C**). This approximately 20% increase in explanatory strength over either the reduced structural model or reduced categorical model alone indicates that there is not direct overlap in the variation explained by the two modeling approaches. We thus decided to extend both modeling perspectives to interrogate other NatE MESA receptor families.

### TNF MESA performance is driven by similar structural features identified in VEGF MESA analysis

Following the approach applied to probe VEGF MESA, we next investigated TNF MESA. TNFRs form assemblies prior to ligand binding^37^, and they signal as ligand-bound trimers upon binding the homotrimeric ligand TNF^38,39^, which is a distinct oligomerization state from VEGFRs (which signal as dimers). Notably, TNF MESA receptors exhibited poor performance (but were still ligand-inducible) compared to VEGF MESA in the prior empirical dataset^26^, and we sought to investigate whether similar rules might apply to such differently performing receptors. Given that multiple distinct receptor-ligand complexes could plausibly form in systems using both the TNFR1 and TNFR2 ECDs (e.g., one TNFR1 ECD + two TNFR2 ECDs; two TNFR1 ECDs, + one TNFR2 ECD; three TNFR1 ECDs; etc.), we elected to model the simplest heteromeric scenario, in which one TNFR1 ECD and one TNFR2 ECD bound to a single TNF ligand to quantify ECD features (**Fig. 4A**). For consistency, we generated homomeric structural models of TNFR1 and TNFR2 bound to TNF as homodimers to use (**Fig. 4A**). Further, we did not discern appreciable structural differences as result of ligand type (i.e., membrane-bound proTNF or soluble TNF with different signal peptides, data not shown), the same structural model measurements were applied to the empirical data corresponding to all ligands. Examining the predicted structure, we first noted that the TNFR2 stalk region, which connects the ligand binding region to the TMD, had low pLDDT scores and folded upwards (**Fig. 4A, Supplementary Fig. 7B, C**). This region was not included in previous crystallographic investigations (PDB ID: 3ALQ)^40^, although it has been proposed that the proline-rich stalk is rigid and may serve to preclude the formation of pre-ligand assemblies^41^. The divergence of the TNFR2 stalk regions in the predicted structures, while not physically realistic, concords with the hypothesis that these regions do not closely associate. Since no ECD contacts were detected in the TNFR-TNF predicted structural models (**Fig. 4A**), we evaluated five structural features for this receptor family. We selected log-transformed OS as the dependent variable for TNF MESA analyses to reduce the influence of outliers on the regression models, as the OS distribution was right-skewed (**Supplementary Fig. 8B**). We found moderately strong relationships between each of the four TMD features and log-transformed OS, while ECD distance had a relatively weak relationship with log-transformed OS (**Fig. 4B**). Consistent with VEGF MESA findings, both ECD distance and TMD distance correlated negatively with log-transformed OS, and the number of TMD contacts correlated positively with log-transformed OS. TMD exit angle exhibited the strongest relationship with OS (r = 0.75), indicating that TNF MESAs containing TMDs with ends facing further apart (closer to 180°) tended to have higher log-transformed OS (**Fig. 4B**). Similar to VEGF MESA, TMD distance, TMD contacts, and TMD crossing angle exhibited strong relationships with one another (**Fig. 4C**).

**Figure 4:**
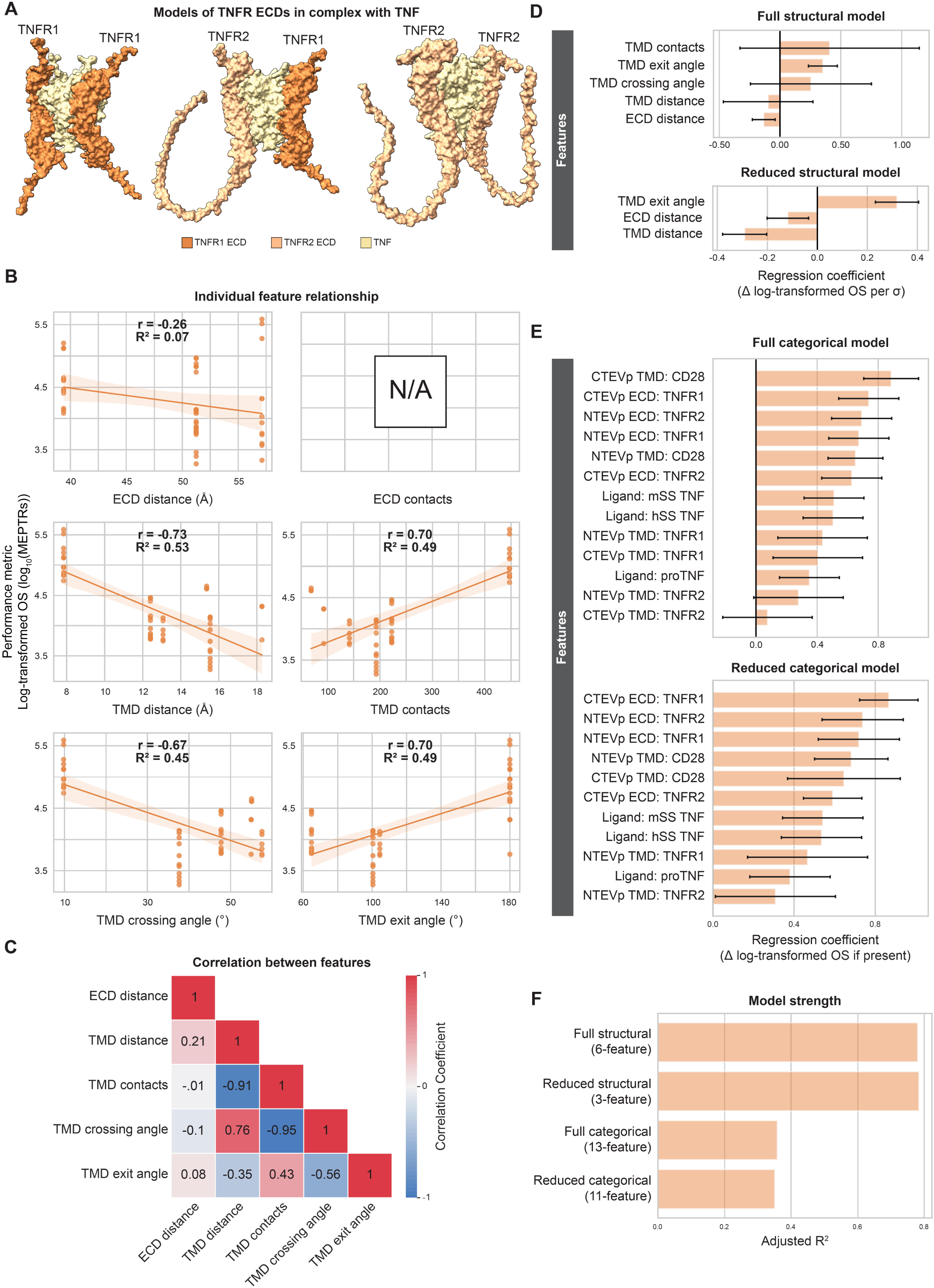
Structural variables explain considerable performance variation in TNF MESA. (**A**) ColabFold-generated models of TNFR ECDs in complex with TNF (tan): TNFR1-TNFR1 (left), TNFR1-TNFR2 (middle), TNFR2-TNFR2 (right) (**Supplementary Data 1, Structures 10-12**). (**B**) Individual structural feature contributions to OS. (**C**) Evaluation of feature independence (Pearson’s correlation coefficients). (**D**) Magnitude of the coefficients for the full (five-feature) and reduced (three- feature) structural models. Structural feature data were normalized to mean = 0 and standard deviation = 1 prior to analyses. Error bars represent +/- 95% confidence intervals calculated using Student’s t- distribution. (**E**) Magnitude of the coefficients for the full (thirteen-feature) and reduced (eleven-feature) categorical models. Error bars represent +/- 95% confidence intervals calculated using Student’s t- distribution. (**F**) Comparison of the four models’ strengths in explaining variation in log-transformed OS, as determined by adjusted R^2^. Abbreviations: tumor necrosis factor receptor (TNFR), tumor necrosis factor receptor 1 (TNFR1), tumor necrosis factor receptor 2 (TNFR2), tumor necrosis factor (TNF), adjusted R^2^ (Adj. R^2^), Pearson’s correlation coefficient (r), coefficient of determination (R^2^), on-state reporter signal (OS), molecules of equivalent PE-TexasRed (MEPTRs), ectodomain (ECD), transmembrane domain (TMD), cluster of differentiation 28 (CD28), N-terminal half of Tobacco Etch Virus Protease (NTEVp), C- terminal half of Tobacco Etch Virus Protease (CTEVp), change in (Δ), standard deviation (σ).

We next explored how combinations of structural or categorical features might explain variation in TNF MESA performance (**Fig. 4D-F**). The full structural model (five features) accounted for much variation in log-transformed OS (Adj. R^2^ = 0.83) (**Fig. 4D**). When this model was reduced as described for VEGF MESA, it retained three terms—ECD distance, TMD contacts, and TMD exit angle—and preserved its explanatory strength (Adj. R^2^ = 0.83) (**Fig. 4F**). Both reduced structural models for VEGF MESA and TNF MESA included terms for ECD distance and TMD contacts, bolstering the importance of these features across NatE MESA receptor families. In contrast, the full categorical model (thirteen, one-hot encoded variables) explained relatively little variation in performance (Adj. R^2^ = 0.36) (**Fig. 4E, F**). Altogether, the structural modeling approach best explained variation in TNF MESA performance and supported conclusions drawn from structural analysis of a distinct MESA family.

### Structural features account for little variation in heterotetrameric receptor performance

We next extended our analysis to receptors whose natural variants form higher order complexes—IL-10 MESA and TGFβ MESA. Although IL-10Rs and TGFβRs derive from different receptor families, they are understood to form receptor-ligand complexes via similar mechanisms. For IL-10Rs, the high-affinity IL-10Rα chain binds on either side of the homodimeric IL-10 ligand, and subsequently the low-affinity IL-10Rβ receptor binds to form a heterotetramer.^42^ Similarly, high-affinity TGFβR2 receptors bind on either side of a homodimeric TGFβ ligand, with the low-affinity TGFβR1 receptors subsequently binding to form a heterotetrameric complex.^43,44^ To evaluate such receptor complexes, we generated structures as heterotetrameric complexes (four receptor ECDs plus the dimeric ligand) (**Fig. 5A, 6A**) and quantified ECD features as the average of the measurements between the heterodimeric ECDs closest to one another. To quantify features for homomeric ECD pairings, we input the sequence to ColabFold as a homodimeric complex (two receptor ECDs plus the dimeric ligand) (**Fig. 5A, 6A**), as each ligand (dimer) contains two binding sites for each receptor. Since the empirical datasets describing performance of each of these MESA receptors^26^ exhibited right-skewed OS distributions, we added a preprocessing step to log-transform OS values to remove the influence of outliers on these regression models (**Supplementary Fig. 8C, D**). Notably, IL-10 MESA exhibited good performance (in terms of ligand-mediated inducibility) while TGFβ MESA variants were generally not very inducible, but since both families exhibited signaling in the presence of ligand, we still investigated use of the log-transformed OS metric for comparing these two receptor families.

**Figure 5:**
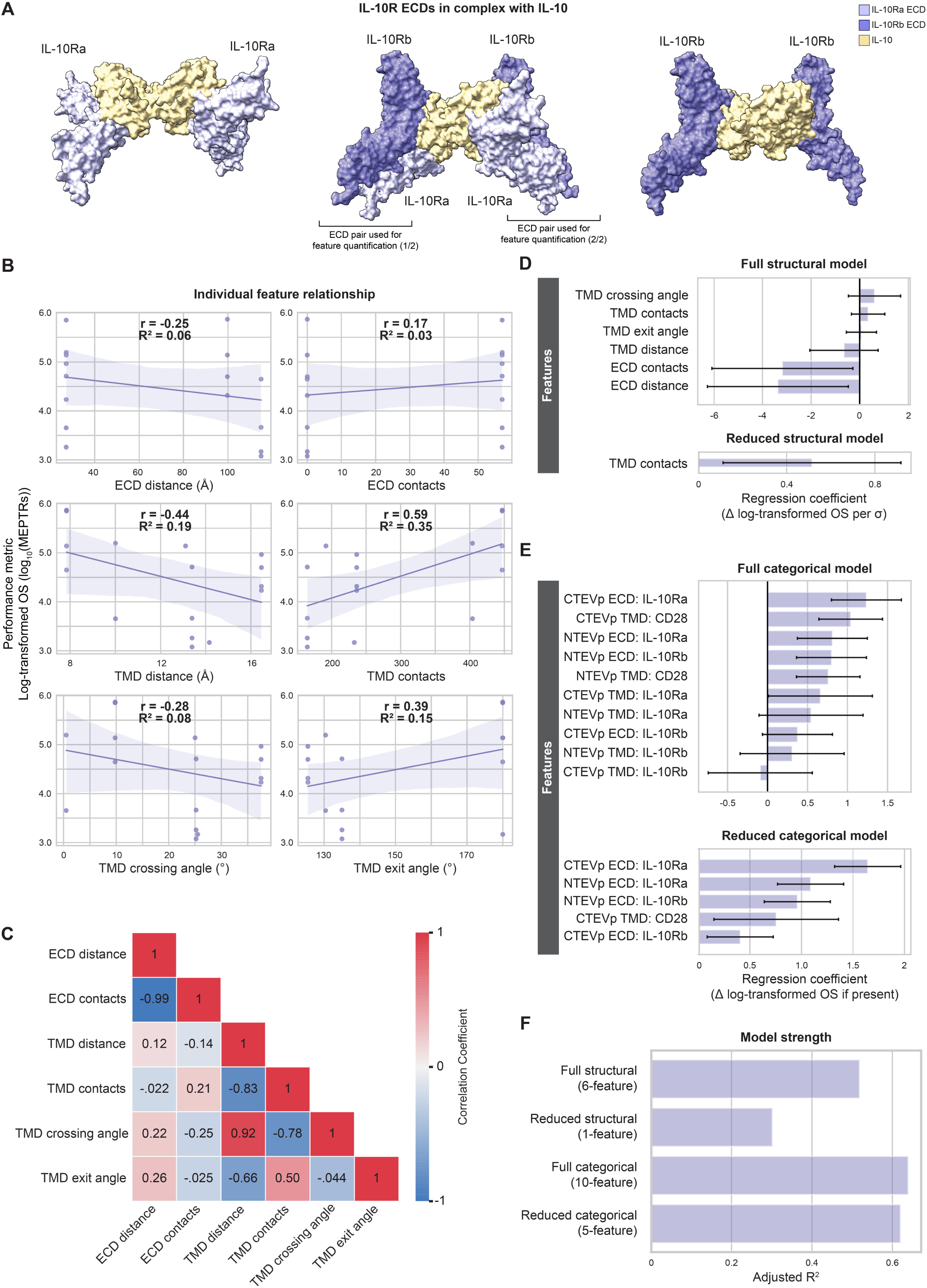
Structural variables explain little performance variation in IL-10 MESA. (**A**) ColabFold-generated models of IL-10R ECDs in complex with IL-10 (tan): IL-10Ra-IL-10Ra (left), IL- 10Ra-IL-10Rb tetramer (middle), IL-10Rb-IL-10Rb (right) (**Supplementary Data 1, Structures 13-15**). Brackets beneath the IL-10Ra-IL-10Rb tetramer (middle) denote ECD pairings that were selected for ECD feature quantification, averaging the measurements between the two pairings. (**B**) Individual structural feature contributions to OS. (**C**) Evaluation of feature independence (Pearson’s correlation coefficients). (**D**) Magnitude of the coefficients for the full (six-feature) and reduced (one-feature) structural models. Structural feature data were normalized to mean = 0 and standard deviation = 1 prior to analyses. Error bars represent +/- 95% confidence intervals calculated using Student’s t-distribution. (**E**) Magnitude of the coefficients for the full (ten-feature) and reduced (five-feature) categorical models. Error bars represent +/- 95% confidence intervals calculated using Student’s t-distribution. (**F**) Comparison of the four models’ strength in explaining variation in log-transformed OS, as determined by adjusted R^2^. Abbreviations: interleukin 10 receptor (IL-10R), interleukin 10 receptor a (IL-10Ra), interleukin 10 receptor b (IL-10Rb), interleukin 10 (IL-10), adjusted R^2^ (Adj. R^2^), Pearson’s correlation coefficient (r), coefficient of determination (R^2^), on-state reporter signal (OS), molecules of equivalent PE-TexasRed (MEPTRs), ectodomain (ECD), transmembrane domain (TMD), cluster of differentiation 28 (CD28), N-terminal half of Tobacco Etch Virus Protease (NTEVp), C-terminal half of Tobacco Etch Virus Protease (CTEVp), change in (Δ), standard deviation (σ).

**Figure 6:**
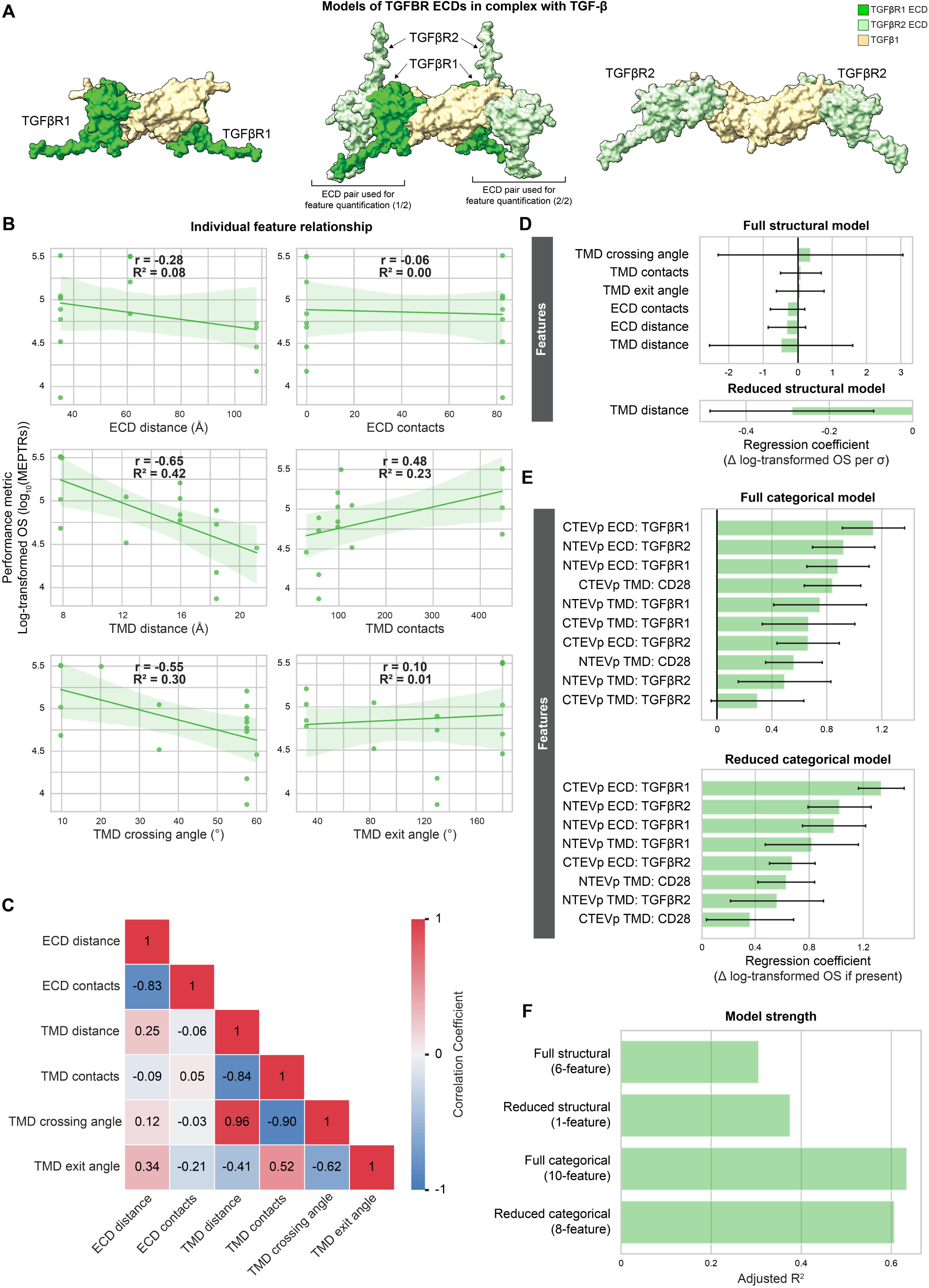
Structural variables explain little performance variation in TGFβ MESA. (**A**) ColabFold-generated models of TGFβR ECDs in complex with IL-10 (tan): TGFβR1-TGFβR1 (left TGFβR1-TGFβR2 tetramer (middle), TGFβR2-TGFβR2 (right) (**Supplementary Data 1, Structures 16- 18**). Brackets beneath the TGFβR1-TGFβR2 tetramer (middle) denote ECD pairings that were selected for ECD feature quantification, averaging the measurements between the two pairings. (**B**) Individual structural feature contributions to OS. (**C**) Evaluation of feature independence (Pearson’s correlation coefficients). (**D**) Magnitude of the coefficients for the full (six-feature) and reduced (one-feature) structural models. Structural feature data were normalized to mean = 0 and standard deviation = 1 prior to analyses. Error bars represent +/- 95% confidence intervals calculated using Student’s t-distribution. (**E**) Magnitude of the coefficients for the full (ten-feature) and reduced (eight-feature) categorical models. Error bars represent +/- 95% confidence intervals calculated using Student’s t-distribution. (**F**) Comparison of the four models’ strength in explaining variation in log-transformed OS, as determined by adjusted R^2^. Abbreviations: transforming growth factor β receptor (TGFβR), transforming growth factor β receptor 1 (TGFβR1), transforming growth factor β receptor 2 (TGFβR2), transforming growth factor β (TGFβ), adjusted R^2^ (Adj. R^2^), Pearson’s correlation coefficient (r), coefficient of determination (R^2^), on-state reporter signal (OS), molecules of equivalent PE-TexasRed (MEPTRs), ectodomain (ECD), transmembrane domain (TMD), cluster of differentiation 28 (CD28), N-terminal half of Tobacco Etch Virus Protease (NTEVp), C-terminal half of Tobacco Etch Virus Protease (CTEVp), change in (Δ), standard deviation (σ).

We first investigated the ligand-inducible, IL-10 MESA family (**Fig. 5A**). Structural features generally explained little variation in performance metrics for IL-10 MESA as compared to VEGF MESA and TNF MESA (**Fig. 5B**). The four TMD features had greater explanatory strength than did ECD features, with TMD contacts contributing the most information (R^2^ = 0.35), and some TMD features again encoded related information (**Fig. 5C**). A full linearly combined structural model accounted for over half of the variation in OS (Adj. R^2^ = 0.52) (**Fig. 5D, F**), but model reduction eliminated all features except for TMD contacts (**Fig. 5D**) and yielded little explanatory strength (Adj. R^2^ = 0.30). The full categorical model (including ten one-hot encoded variables) explained substantial variation in OS (Adj. R^2^ = 0.64) (**Fig. 5E, F**), and the reduced (five-feature) model still explained much variation (Adj. R^2^ = 0.62) (**Fig. 5E, F**). Since this relative inadequacy of the structural model was surprising, we next investigated whether these trends held for the mechanistically similar TGFβ MESA.

Our analysis of TGFβ MESA yielded observations mirroring those observed with IL-10 MESA (**Fig. 6A**). Again, TMD features yielded the strongest relationships with log-transformed OS (**Fig. 6B**), and information contained in such features was related (**Fig. 6C**). Both the full (six-feature) structural model and the reduced model (one term: TMD distance) explained little variation in performance (**Fig. 6D, F**). Again, the full (ten-feature) categorical model and its reduced (eight-feature) form explained a considerably greater proportion of the variation in performance than did the structural feature models (**Fig. 6E, F**).

The combined analysis of these heterotetrameric receptors suggests clues that could indicate limitations to our current structure prediction-guided analysis. One possible explanation for the low explanatory strength of structural feature models for IL-10 MESA and TGFβ MESA is that heterotetrameric receptors may form higher-order complexes that are not reflected by our models. For example, IL-10Rα NTEVp paired with IL-10Rα CTEVp is predicted to have ∼100Å distance between ECDs when bound to IL-10, and that distance is far too wide to allow TMD interactions or TEVp reconstitution (**Supplementary Fig. 2D**), yet this MESA combination was empirically observed to exhibit ligand-induced signaling.^26^ As a next step, these findings motivated a systematic evaluation of conclusions across the ensemble of NatE MESA families investigated.

### Synthesizing general drivers of MESA performance across receptor families

Finally, given the various relationships discovered between predicted structure and function within individual NatE MESA receptor families, we investigated whether there exist general principles that apply across receptor families. First, we noted that the correlation coefficients associated with each structural feature exhibited consistent signs across each of the individual receptor family models (**Fig. 7A**). The consistency of these relationships supports our hypothesis that global trends exist between structural features and MESA performance, even across various categories of receptor-ligand signaling complexes (dimeric, trimeric, tetrameric). We next combined the four NatE MESA receptor families into a single dataset (“all receptor”) to explore the relationship between each feature and log-transformed OS. Of the six structural features, TMD distance explained the greatest amount of variation in OS of any individual feature (R^2^ = 0.32) (**Supplementary Fig. 1B**). The full structural model (six features) explains some variation (Adj. R^2^ = 0.32), and reduction of this model (retaining three features: ECD contacts, TMD distance, and TMD exit angle) modestly improved the explanatory strength of the model (Adj. R^2^ = 0.44) (**Fig. 7B**). It is unsurprising that such an approach is unable to capture all the variation in this combined set, but the comparison does highlight some intriguing differences among the receptor designs. For example, by comparing the observed vs. predicted log-transformed OS for each receptor (**Fig. 7C**), we see that the subset of VEGF MESA receptors with highest log-transformed OS yields the tightest correlation between predicted and observed performance, and within the TNF MESA dataset, there exist many designs which differ in observed performance but are not distinguished by structural features retained in the reduced model. We also assessed whether structural features could explain variation within a distinct performance metric, FI (log-transformed). Individually, the four TMD features explained very little variation, but ECD contacts explained substantial variation in log-transformed FI (Adj. R^2^ = 0.52) (**Supplementary Fig. 1D**). In a multi-feature model, including both full (six feature) and reduced (four feature) forms, similar trends were observed (**Fig. 7D, E**). One possible interpretation is that strong receptor-ligand complexes, bolstered by interactions between the two chains, are conducive to high FI.

**Figure 7:**
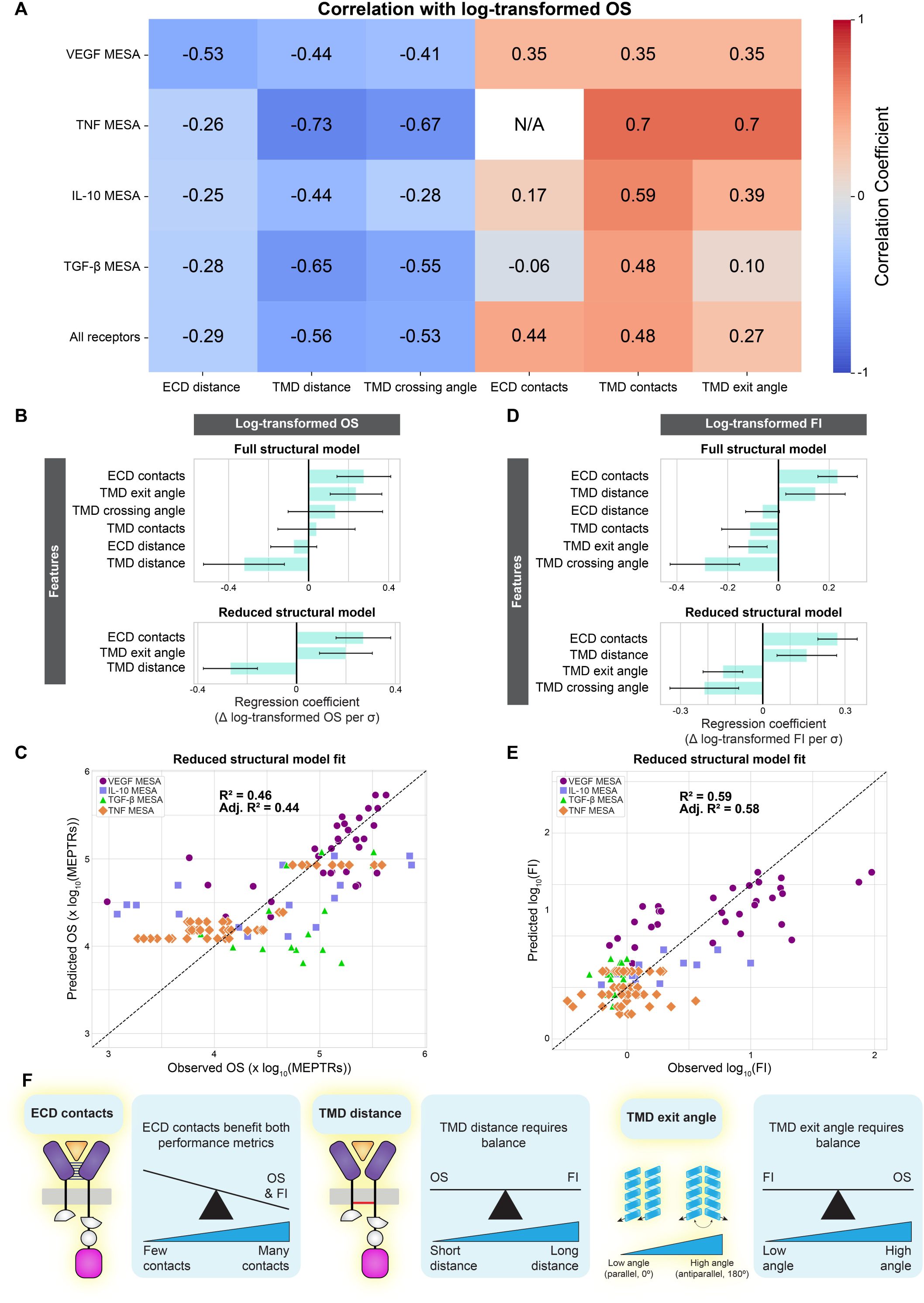
Structure-function relationships across NatE MESA receptor families. (**A**) Comparison of structural feature contributions to performance across receptor families (Pearson’s correlation coefficient between each feature and log-transformed OS). “All receptors” considers all data from the four distinct receptors families as a single dataset. In all but one of the 29 relationships (TGFβR, TMD contacts), the direction of the relationship (positive or negative) is consistent across all five datasets. (**B**) Magnitude of the coefficients for the full (six-feature) and reduced (three-feature) structural models explaining variation in log-transformed OS. Structural feature data were normalized to mean = 0 and standard deviation = 1 prior to performing analyses. Error bars represent +/- 95% confidence intervals calculated using Student’s t-distribution. (**C**) The reduced structural model fit, explaining variation in log- transformed OS. The x=y line would represent a perfect fit. (**D**) Magnitude of the coefficients for the full (six-feature) and reduced (four-feature) structural model explaining variation in log-transformed FI. Structural feature data were normalized to mean = 0 and standard deviation = 1 prior to analyses. Error bars represent +/- 95% confidence intervals calculated using Student’s t-distribution. (**E**) The reduced structural model fit, explaining variation in log-transformed FI. (**F**) A synthesis of general structure-function relationships observed across MESA receptor families. Abbreviations: on-state reporter signal (OS), vascular endothelial growth factor receptor (VEGFR), tumor necrosis factor receptor (TNFR), interleukin 10 receptor (IL-10R), transforming growth factor β receptor (TGFβR), modular extracellular sensor architecture (MESA), ectodomain (ECD), transmembrane domain (TMD), adjusted R^2^ (Adj. R^2^), Pearson’s correlation coefficient (r), coefficient of determination (R^2^), molecules of equivalent PE-TexasRed (MEPTRs), fold induction (FI), change in (Δ), standard deviation (σ).

Altogether, this comparative analysis complements our investigation of individual receptor families to suggest potential design rules that may guide future receptor engineering (**Fig. 7F**). For example, increasing of ECD contacts correlated with improved OS and FI across receptor families, suggesting that this property could be used to select or engineer protein ectodomains to build or optimize receptors. However, both TMD distance and TMD exit angle impact OS and FI differently, suggesting that one might select these properties to advance the performance metric of greatest interest, or alternatively, build a library of receptor variants sampling these metrics to increase the probability of obtaining a desirable balance. In general, the fact that both our individual receptor investigations and our combined analysis yielded experimentally testable hypotheses underscores the value of the predicted structure-guided synthetic receptor analysis pioneered here.

## DISCUSSION

This study explored whether predicted protein structures might identify features that can help understand determinants of performance for a model synthetic receptor class, and the outcomes indicate that this approach has promise. While we encountered some limitations when attempting to generate structures of complete chimeric engineered receptors, generating separate models of the ECD and the TMD domains still enabled measuring structural variables of interest. Even a relatively simple set of features accounted for substantial performance variation for VEGF and TNF MESAs, and such relationships tended to be weaker with the heterotetrameric IL-10 MESA and TGFβ MESA. Across the four families of NatE MESAs, the signs of the relationships between each feature and OS were largely conserved. Shorter ECD and TMD distances correlated with higher OS, as did greater numbers of ECD and TMD contacts, which agree with our general model of MESA signaling, in that shorter distances and stronger receptor-ligand complexes tend to benefit performance^35^. Within individual receptor family analyses, ECD distance and ECD contacts tended to have strong collinearity, as did TMD distance and TMD contacts. This is not necessarily surprising as there is an inherent relationship between distances and interchain interactions. TMD crossing angle tended to strongly correlate with TMD distance, although it often explained less variation in either OS or FI. Through TMD exit angle, we attempted to capture some aspect of the relative orientation of NTEVp and CTEVp when receptor-ligand complexes form, as we surmise that this is a key factor in TEVp reconstitution propensity and subsequent signaling. While TMD exit angle played a substantial role in the TNF MESA and “all receptor” models, interpretation of its role is less straightforward (e.g., compared to contact and distance measurements) as it only indirectly relates to NTEVp and CTEVp orientation. This case highlights a general challenge in that we might expect structural model predictions to be more accurate for designs in which the receptor functions (i.e., when TEVp fragments reconstitute upon ligand binding) than for designs in which structural constraints preclude formation of the viable signaling state (and the ensemble of “frustrated” structures may be more difficult to predict). Although these constraints guided our selection of simple structural features, we nonetheless gleaned substantial insights. Overall, this approach identified correlations between structural features and MESA performance metrics, and this opens an exciting future avenue to experimentally interrogate which, if any, of these correlations are causal or predictive.

Our post hoc analyses enabled us to pose new, testable hypotheses as to how one might guide or improve MESA performance through protein engineering, and here we contemplate an example. TNF MESAs exhibited relatively poor performance in the empirical NatE MESA investigation^26^, lacking a useful FI, even after evaluating changes such as varying split-TEVp mutants to reduce background signal. This study suggests a distinct opportunity—addressing the lack of any contacts between TNFR chains, as ECD contacts have a moderate relationship with higher OS and a strong relationship with higher FI. A contributing factor to the lack of ECD contacts could be the TNFR stalk region, which connects the ligand-binding domain to the TMD. The TNFR2 stalk region is reported to have repulsive interchain forces, which reduces natural TNFR2 responsiveness to soluble TNF.^41,45^ In our work, both the TNFR1 and TNFR2 stalk regions were predicted to lack secondary structure, though the confidence scores in these regions are very low (**Supplementary Fig. 7A-C**). A plausible improvement could be replacing the natural stalk regions with a contact-rich synthetic linker domain pair. Moreover, one might select these linkers to sample a range of interaction propensities (to avoid driving ligand-independent interactions), while focusing on the pair having a relatively short ECD distance. Exploring a set of linkers with similar structure, but varying amounts of interchain contacts, could help evaluate whether the relationship between ECD contacts and FI is causal.

The selection of TMD pairs has a large influence on MESA performance^35^, which resonates with findings observed with other synthetic receptors^36,46^, and our study guides consideration of this feature for receptor engineering. We found that TMDs with greater numbers of contacts and closer C-termini tended to elicit higher OS. However, neither of these features correlated appreciably with FI (using the linear regression evaluated). One feature not quantified in this investigation, but which may influence FI, is the propensity of TMDs to aggregate. In general, we speculate that TMD aggregation leads to high amounts of background signaling, thereby reducing FI. We and others have posited that since CD28 proteins form clusters to signal, use of the CD28 TMD may drive high background signaling.^35,36^ Altogether, optimizing receptor performance in a manner that balances such outcomes would benefit from detailed models as to how TMD sequence impacts all of these biophysical properties. A promising avenue of future work may be to combine experimental characterization of TMD interactions, and statistical (machine learning) or biophysical models trained on such data, with structural models incorporating both TMDs and intracellular and extracellular domains to identify actionable design principles relevant to this important receptor domain.

This investigation identified multiple future opportunities for improving or extending protein structure prediction methods. A critical piece of information that we were unable to directly capture in this investigation is the spatial relationship and orientation between the NTEVp and CTEVp within receptor-ligand complexes. In our hands, ColabFold could not consistently produce models with NTEVp and CTEVp folded—even for receptors whose experimental performance suggests reconstitution occurs, which appeared to be related to a lack of sequence coverage in the MSA. A key clue was that receptors with shorter ECDs tended to obtain adequate coverage of the TEVp regions in the MSA, often resulting in a fully reconstituted TEVp. Thus, one possible solution could include adjusting the ColabFold algorithm to more specifically accommodate the particular needs for modeling chimeric proteins. For example, sequence alignment queries may be modified to ensure coverage over all components by allowing users to define domains within a single chain and adapting the MSA search and construction process to each modeling goal. However, even in cases where a folded TEVp is predicted, a note of caution is warranted—since the TMD (and the juxtamembrane region) often received very low confidence scores, it is possible that ColabFold prioritizes score-benefitting reconstitution of the TEVp over prediction of the TMD and juxtamembrane domains. This issue also reflects the aforementioned challenge that we expect structure prediction to be most accurate for receptors that do successfully form a signaling-competent complex, while it would remain more challenging to generate or interpret structural predictions for “frustrated” receptors that signal poorly or not at all. An additional opportunity identified in our investigation of heterotetrameric receptors is the potential need to consider higher order receptor complexes. Beyond the computational challenge associated with predicting such structures, it will be important to consider how to best evaluate the ensemble of states and complexes that might be relevant drivers of receptor function. Finally, an additional opportunity is the use of detailed biophysical models, such as molecular dynamics simulations, to directly test putative causality and drivers of performance including those identified in this study.

### The Bigger Picture

We expect increased use of structural modeling to analyze and guide receptor engineering as these tools continue to improve. The present study comprises a proof-of-concept that high information content synthetic receptor libraries can be experimentally characterized and analyzed using structure prediction-enabled machine learning methods to guide subsequent refinement, as discussed above. A natural evolution of this workflow is to generate libraries that explore greater design space and ultimately automated design of subsequent library generations to accelerate iterative optimization, as supportive methods such as BioAutoMATED^47^ are increasingly available and accessible. Ultimately, structural predictions may become indispensable tools for improving our understanding of structure-function relationships in engineered receptors, accelerating the development of high-performing parts that enable diverse applications in biotechnology and medicine.

## CONCLUSION

Protein structure prediction tools are a powerful emerging technology that can be leveraged to generate insights into structural contributions to engineered receptor function. In this investigation, we piloted and validated this approach using MESA receptors as a case study, and we expect that similar methodology could be applied to many other engineered receptors. We identified a core set of structural features that exhibited consistent correlations with receptor performance across four different families of MESAs derived from diverse cytokine receptors. These analyses enabled us to pose specific, testable hypotheses as to how to improve MESA performance, which is an exciting avenue for future work. This proof of principle establishes the feasibility of employing structure prediction to evaluate and guide synthetic receptor design.

## METHODS

### Generation of ColabFold Models

The publicly available Google Colaboratory notebook, ColabFold, version 1.5.2 (or newer), was used to create AlphaFold models.^28^ We used the following parameter settings: num_relax = 1, template_mode = pdb100, msa_mode = mmseqs2_uniref_env, pair_mode = unpaired_paired, model_type = auto (alphafold2_multimer_v3), num_recycles = auto, relax_max_iterations = 200. To quantify features of each ectodomain model, we selected the top-ranking model of the five generated, which was relaxed using the automatic amber relax protocol in the Colab notebook. Each amino acid sequence analyzed in this study is enumerated and listed in **Supplementary Table 2**.

### Generation of PREDDIMER Models

To model TMD dimers, we used PREDDIMER.^30,31^ For homodimeric TMD pairings, we selected the top-ranking model based on PREDDIMER’s F_scor_, which accounts for the packing quality of each dimer, from which to quantify features. For heterodimeric TMD pairings, we noted that results differed as a function of which sequence was entered as the first or second in the pair. To account for these differences, we generated two sets of models for each heterodimeric pair, with each of the two TMDs being entered as the first sequence. We then quantified the relevant features from each of the top-ranking models and calculated the arithmetic mean metric for use in subsequent analyses. Each amino acid sequence analyzed is listed in **Supplementary Table 2**, and F_scor_ scores for each dimer are listed in **Supplementary Table 3**.

### Quantification of features

To quantify features of the ECD models, we used ChimeraX-1.6.1.^48^ To determine the ECD C-terminal distance, we selected the final residue in each chain that was predicted to be a part of a secondary structural element (e.g., α helix or β sheet) and measured the distance between the Cα of each residue using the Distance tool. The number of contacts between each chain was quantified using ChimeraX’s Contacts tool, using VDW overlap tolerance = 0.40Å and ignored interactions between 4 or fewer bonds apart, which are the default settings. To detect only inter-chain interactions, we selected one chain using the select tool and specified the Contacts tool to limit by selection between atoms selected in the other chain. The number of contacts between each TMD in the PREDDIMER models was quantified using ChimeraX’s Contacts tool, using VDW overlap ≥ 0.40Å, ignoring interactions between atoms 4 or fewer bonds apart, and limiting by selection to only interactions between atoms of the two different TMD chains. The TMD C-terminal distance was quantified by measuring the distance between the Cα atoms of the final residue between the two chains. The TMD chi-angle was quantified as calculated by PREDDIMER.

The relative TMD exit angle is a metric which we created. To create this measurement, we extracted the X and Y coordinates of both the Cα and C (carboxyl carbon) to create a vector between these two points. These data were acquired for each chain, creating two vectors, from Cα and C for each chain. We then used the dot product and the magnitudes of these two vectors to calculate the cosine, which was converted to an angle in the unit of degrees.

### Statistical Analyses

Regression analyses were conducted in Python using statsmodels 0.14.2, using the Ordinary Least Squares method. For both simple linear regression (single feature) and multiple linear regression models, a fit constant term (y-intercept) was employed, allowing the model to better fit the data by not forcing the regression line through the origin, as that constraint would have no physical meaning. For multiple linear regression models, the values for each structural feature (independent variables) were standardized to a mean = 0 and standard deviation = 1 prior to analysis. To reduce the multiple linear regression models, we first determined the variance inflation factor (VIF) for each feature, and removed the feature with the highest VIF, using the cutoff VIF ≥ 10. We subsequently calculated VIF values for the remaining features and removed the largest value (if ≥ 10) until all VIF scores were below 10. Finally, we performed backward elimination, iteratively removing the feature with the largest p-value until all features were statistically significant at α = 0.05.

## ACKNOWLEDGEMENTS

Molecular graphics and analyses performed with UCSF ChimeraX, developed by the Resource for Biocomputing, Visualization, and Informatics at the University of California, San Francisco, with support from National Institutes of Health R01-GM129325 and the Office of Cyber Infrastructure and Computational Biology, National Institute of Allergy and Infectious Diseases.

## AUTHOR CONTRIBUTIONS

W.K.C: conceptualization, methodology, data curation, formal analysis, investigation, resources, software, visualization, writing – original draft, writing – review & editing. A.C.: methodology, data curation, writing – review & editing. H.I.E: writing – review & editing. J.N.L.: conceptualization, funding acquisition, project administration, supervision, writing – review and editing. Manuscript was reviewed and edited by all authors prior to submission.

## SUPPLEMENTARY MATERIAL

Supplementary material online includes **Supplementary Information** (**Supplementary Figures 1-9**, **Supplementary Tables 1, 2, 3**), **Supplementary Data 1** (archive containing ColabFold-generated structural models used in this study) and **Supplementary Data 2** (archive containing PREDDIMER- generated TMD dimers used in this study)^49^, and a **Source Data** file containing values used to generate all plots reported.

## REFERENCES

1. June CH, O’Connor RS, Kawalekar OU, et al. CAR T cell immunotherapy for human cancer. Science 2018;359(6382):1361–1365, doi:10.1126/science.aar6711

2. Mitra A, Barua A, Huang L, et al. From bench to bedside: the history and progress of CAR T cell therapy. Frontiers in Immunology 2023;14(doi:10.3389/fimmu.2023.1188049

3. Chung JB, Brudno JN, Borie D, Kochenderfer JN. Chimeric antigen receptor T cell therapy for autoimmune disease. Nature Reviews Immunology 2024, doi:10.1038/s41577-024-01035-3

4. Blache U, Tretbar S, Koehl U, et al. CAR T cells for treating autoimmune diseases. RMD Open 2023;9(4), doi:10.1136/rmdopen-2022-002907

5. Manhas J, Edelstein HI, Leonard JN, Morsut L. The evolution of synthetic receptor systems. Nat Chem Biol 2022;18(3):244–255, doi:10.1038/s41589-021-00926-z

6. Guedan S, Calderon H, Posey AD, Maus MV. Engineering and Design of Chimeric Antigen Receptors. Mol Ther-Meth Clin D 2019;12(145–156, doi:10.1016/j.omtm.2018.12.009

7. Morsut L, Roybal KT, Xiong X, et al. Engineering Customized Cell Sensing and Response Behaviors Using Synthetic Notch Receptors. Cell 2016;164(4):780–791, doi:10.1016/j.cell.2016.01.012

8. Roybal KT, Williams JZ, Morsut L, et al. Engineering T Cells with Customized Therapeutic Response Programs Using Synthetic Notch Receptors. Cell 2016;167(2):419-+, doi:10.1016/j.cell.2016.09.011

9. Zhu I, Liu R, Garcia JM, et al. Modular design of synthetic receptors for programmed gene regulation in cell therapies. Cell 2022;185(8):1431-+, doi:10.1016/j.cell.2022.03.023

10. Scheller L, Strittmatter T, Fuchs D, et al. Generalized extracellular molecule sensor platform for programming cellular behavior. Nat Chem Biol 2018;14(7):723–729, doi:10.1038/s41589-018-0046-z

11. Daringer NM, Dudek RM, Schwarz KA, Leonard JN. Modular extracellular sensor architecture for engineering mammalian cell-based devices. ACS Synth Biol 2014;3(12):892–902, doi:10.1021/sb400128g

12. Schwarz KA, Daringer NM, Dolberg TB, Leonard JN. Rewiring human cellular input-output using modular extracellular sensors. Nat Chem Biol 2017;13(2):202–209, doi:10.1038/nchembio.2253

13. Singh N, Frey NV, Engels B, et al. Antigen-independent activation enhances the efficacy of 4- 1BB-costimulated CD22 CAR T cells. Nat Med 2021;27(5):842–850, doi:10.1038/s41591-021-01326-5

14. Li N, Quan A, Li D, et al. The IgG4 hinge with CD28 transmembrane domain improves VHH-based CAR T cells targeting a membrane-distal epitope of GPC1 in pancreatic cancer. Nature Communications 2023;14(1):1986, doi:10.1038/s41467-023-37616-4

15. Hwang MS, Miller MS, Thirawatananond P, et al. Structural engineering of chimeric antigen receptors targeting HLA-restricted neoantigens. Nature Communications 2021;12(1):5271, doi:10.1038/s41467-021-25605-4

16. Bugge K, Lindorff-Larsen K, Kragelund BB. Understanding single-pass transmembrane receptor signaling from a structural viewpoint—what are we missing? The FEBS journal 2016;283(24):4424–4451

17. Cai K, Zhang X, Bai XC. Cryo-electron Microscopic Analysis of Single-Pass Transmembrane Receptors. Chem Rev 2022;122(17):13952–13988, doi:10.1021/acs.chemrev.1c01035

18. Cheung J, Wazir S, Bell DR, et al. Crystal Structure of a Chimeric Antigen Receptor (CAR) scFv Domain Rearrangement Forming a VL-VL Dimer. Crystals 2023;13(4):710

19. NobelPrize.org. They cracked the code for proteins’ amazing structures. NobelPrize.org: Nobel Prize Outreach AB 2024; 2024.

20. Jumper J, Evans R, Pritzel A, et al. Highly accurate protein structure prediction with AlphaFold. Nature 2021;596(7873):583-+, doi:10.1038/s41586-021-03819-2

21. Abramson J, Adler J, Dunger J, et al. Accurate structure prediction of biomolecular interactions with AlphaFold 3. Nature 2024;630(8016):493–500, doi:10.1038/s41586-024-07487-w

22. Evans R, O’Neill M, Pritzel A, et al. Protein complex prediction with AlphaFold-Multimer. bioRxiv 2022;2021.10.04.463034, doi:10.1101/2021.10.04.463034

23. Baek M, DiMaio F, Anishchenko I, et al. Accurate prediction of protein structures and interactions using a three-track neural network. Science 2021;373(6557):871–876, doi:10.1126/science.abj8754

24. Pogozheva ID, Cherepanov S, Park SJ, et al. Structural Modeling of Cytokine-Receptor-JAK2 Signaling Complexes Using AlphaFold Multimer. J Chem Inf Model 2023;63(18):5874–5895, doi:10.1021/acs.jcim.3c00926

25. Chang JF, Landmann JH, Chang TC, et al. Rational protein engineering to enhance MHC-independent T cell receptors. Cancer Discov 2024, doi:10.1158/2159-8290.Cd-23-1393

26. Edelstein HI, Cosio A, Ezekiel ML, et al. Conversion of natural cytokine receptors into orthogonal synthetic biosensors. bioRxiv 2024;2024.03.23.586421, doi:10.1101/2024.03.23.586421

27. Dolberg TB, Meger AT, Boucher JD, et al. Computation-guided optimization of split protein systems. Nat Chem Biol 2021;17(5):531–539, doi:10.1038/s41589-020-00729-8

28. Mirdita M, Schutze K, Moriwaki Y, et al. ColabFold: making protein folding accessible to all. Nat Methods 2022;19(6):679–682, doi:10.1038/s41592-022-01488-1

29. Kim G, Lee S, Levy Karin E, et al. Easy and accurate protein structure prediction using ColabFold. Nat Protoc 2024, doi:10.1038/s41596-024-01060-5

30. Polyansky AA, Chugunov AO, Volynsky PE, et al. PREDDIMER: a web server for prediction of transmembrane helical dimers. Bioinformatics 2014;30(6):889–90, doi:10.1093/bioinformatics/btt645

31. Polyansky AA, Volynsky PE, Efremov RG. Multistate organization of transmembrane helical protein dimers governed by the host membrane. J Am Chem Soc 2012;134(35):14390–400, doi:10.1021/ja303483k

32. Sarabipour S, Ballmer-Hofer K, Hristova K. VEGFR-2 conformational switch in response to ligand binding. Elife 2016;5(e13876, doi:10.7554/eLife.13876

33. Chakraborty MP, Das D, Mondal P, et al. Molecular basis of VEGFR1 autoinhibition at the plasma membrane. Nat Commun 2024;15(1):1346, doi:10.1038/s41467-024-45499-2

34. Yang Y, Xie P, Opatowsky Y, Schlessinger J. Direct contacts between extracellular membrane-proximal domains are required for VEGF receptor activation and cell signaling. P Natl Acad Sci USA 2010;107(5):1906–1911, doi:10.1073/pnas.0914052107

35. Edelstein HI, Donahue PS, Muldoon JJ, et al. Elucidation and refinement of synthetic receptor mechanisms. Synth Biol (Oxf) 2020;5(1):ysaa017, doi:10.1093/synbio/ysaa017

36. Elazar A, Chandler NJ, Davey AS, et al. De novo-designed transmembrane domains tune engineered receptor functions. eLife 2022;11(e75660, doi:10.7554/eLife.75660

37. Chan FK, Chun HJ, Zheng L, et al. A domain in TNF receptors that mediates ligand-independent receptor assembly and signaling. Science 2000;288(5475):2351–4, doi:10.1126/science.288.5475.2351

38. D’Arcy A, Banner DW, Janes W, et al. Crystallization and preliminary crystallographic analysis of a TNF-beta-55 kDa TNF receptor complex. J Mol Biol 1993;229(2):555–7, doi:10.1006/jmbi.1993.1055

39. Wajant H, Siegmund D. TNFR1 and TNFR2 in the Control of the Life and Death Balance of Macrophages. Front Cell Dev Biol 2019;7(91, doi:10.3389/fcell.2019.00091

40. Mukai Y, Nakamura T, Yoshikawa M, et al. Solution of the structure of the TNF-TNFR2 complex. Sci Signal 2010;3(148):ra83, doi:10.1126/scisignal.2000954

41. Richter C, Messerschmidt S, Holeiter G, et al. The tumor necrosis factor receptor stalk regions define responsiveness to soluble versus membrane-bound ligand. Mol Cell Biol 2012;32(13):2515–29, doi:10.1128/mcb.06458-11

42. Kotenko SV, Krause CD, Izotova LS, et al. Identification and functional characterization of a second chain of the interleukin-10 receptor complex. EMBO J 1997;16(19):5894–903, doi:10.1093/emboj/16.19.5894

43. Dore JJ, Jr., Edens M, Garamszegi N, Leof EB. Heteromeric and homomeric transforming growth factor-beta receptors show distinct signaling and endocytic responses in epithelial cells. J Biol Chem 1998;273(48):31770–7, doi:10.1074/jbc.273.48.31770

44. Vander Ark A, Cao J, Li X. TGF-β receptors: In and beyond TGF-β signaling. Cellular Signalling 2018;52(112–120, 10.1016/j.cellsig.2018.09.002

45. Medler J, Kucka K, Wajant H. Tumor Necrosis Factor Receptor 2 (TNFR2): An Emerging Target in Cancer Therapy. Cancers (Basel) 2022;14(11), doi:10.3390/cancers14112603

46. Teng F, Cui T, Zhou L, et al. Programmable synthetic receptors: the next-generation of cell and gene therapies. Signal Transduction and Targeted Therapy 2024;9(1):7

47. Valeri JA, Soenksen LR, Collins KM, et al. BioAutoMATED: An end-to-end automated machine learning tool for explanation and design of biological sequences. Cell systems 2023;14(6):525–542. e9

48. Meng EC, Goddard TD, Pettersen EF, et al. UCSF ChimeraX: Tools for structure building and analysis. Protein Science 2023;32(11):e4792, 10.1002/pro.4792

49. Corcoran WK, Cosio A, Edelstein HI, Leonard JN. Supplementary Data for “Exploring structure-function relationships in engineered receptor performance using computational structure prediction”. Zenodo.org; 2024. doi: 10.5281/zenodo.14050835

