## Supplementary Information for "Exploring structure-function relationships in engineered receptor performance using computational structure prediction"

**Contents:**

Supplementary Figures 1-9
Supplementary Tables 1-3

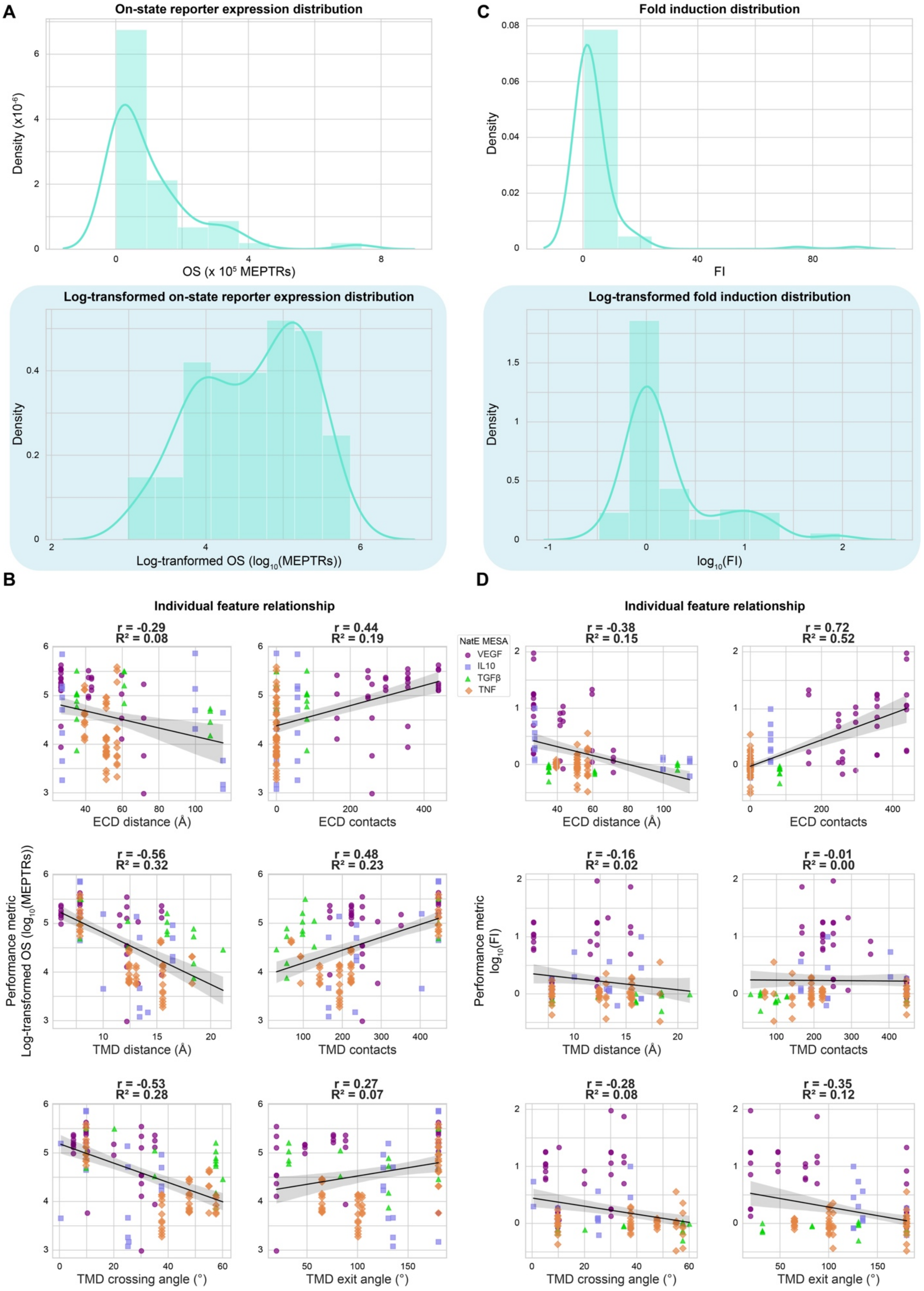
A
On-state reporter expression distribution
Density (x10^-6)
OS (x 10^5 MEPTRs)
Log-transformed on-state reporter expression distribution
Density
Log-tranformed OS (log10(MEPTRs))
C
Fold induction distribution
Density
FI
Log-transformed fold induction distribution
Density
log10(FI)
B
Individual feature relationship
r = -0.29
R² = 0.08
ECD distance (Å)
r = 0.44
R² = 0.19
ECD contacts
NatE MESA
VEGF
IL10
TGFβ
TNF
Performance metric
Log-transformed OS (log10(MEPTRs))
r = -0.56
R² = 0.32
TMD distance (Å)
r = 0.48
R² = 0.23
TMD contacts
r = -0.53
R² = 0.28
TMD crossing angle (°)
r = 0.27
R² = 0.07
TMD exit angle (°)
D
Individual feature relationship
r = -0.38
R² = 0.15
ECD distance (Å)
r = 0.72
R² = 0.52
ECD contacts
Performance metric
log10(FI)
r = -0.16
R² = 0.02
TMD distance (Å)
r = -0.01
R² = 0.00
TMD contacts
r = -0.28
R² = 0.08
TMD crossing angle (°)
r = -0.35
R² = 0.12
TMD exit angle (°)

**Supplementary Figure 1: Identifying structural features that can explain global trends in MESA performance**

(**A**) All four datasets (VEGF MESA, TNF MESA, IL-10 MESA, TGFβ MESA) were aggregated into a single "all receptor" dataset, with the OS distribution (top) and log-transformed OS (bottom) with fitted kernel density estimate (KDE). Log-transformed OS was selected as the performance metric for further analysis as the OS distribution is right-skewed. (**B**) Evaluation of the linear relationship between each individual structural feature and log-transformed OS. (**C**) The "all receptor" FI distribution (top) and log-transformed FI distribution (bottom) with fitted KDEs. Log-transformed FI was selected as the performance metric for further analysis as the OS distribution has a greater right skew. (**D**) Evaluation of the linear relationship between each individual structural feature and log-transformed FI. Abbreviations: on-state reporter expression (OS), fold induction (FI), kernel density estimate (KDE), molecules of equivalent PE-TexasRed (MEPTRs), adjusted $R^2$ (Adj. $R^2$), Pearson's correlation coefficient (r), ectodomain (ECD), transmembrane domain (TMD), natural ectodomain modular extracellular sensor architecture (NatE MESA), vascular endothelial growth factor (VEGF), interleukin 10 (IL-10), transforming growth factor β (TGFβ), tumor necrosis factor (TNF).

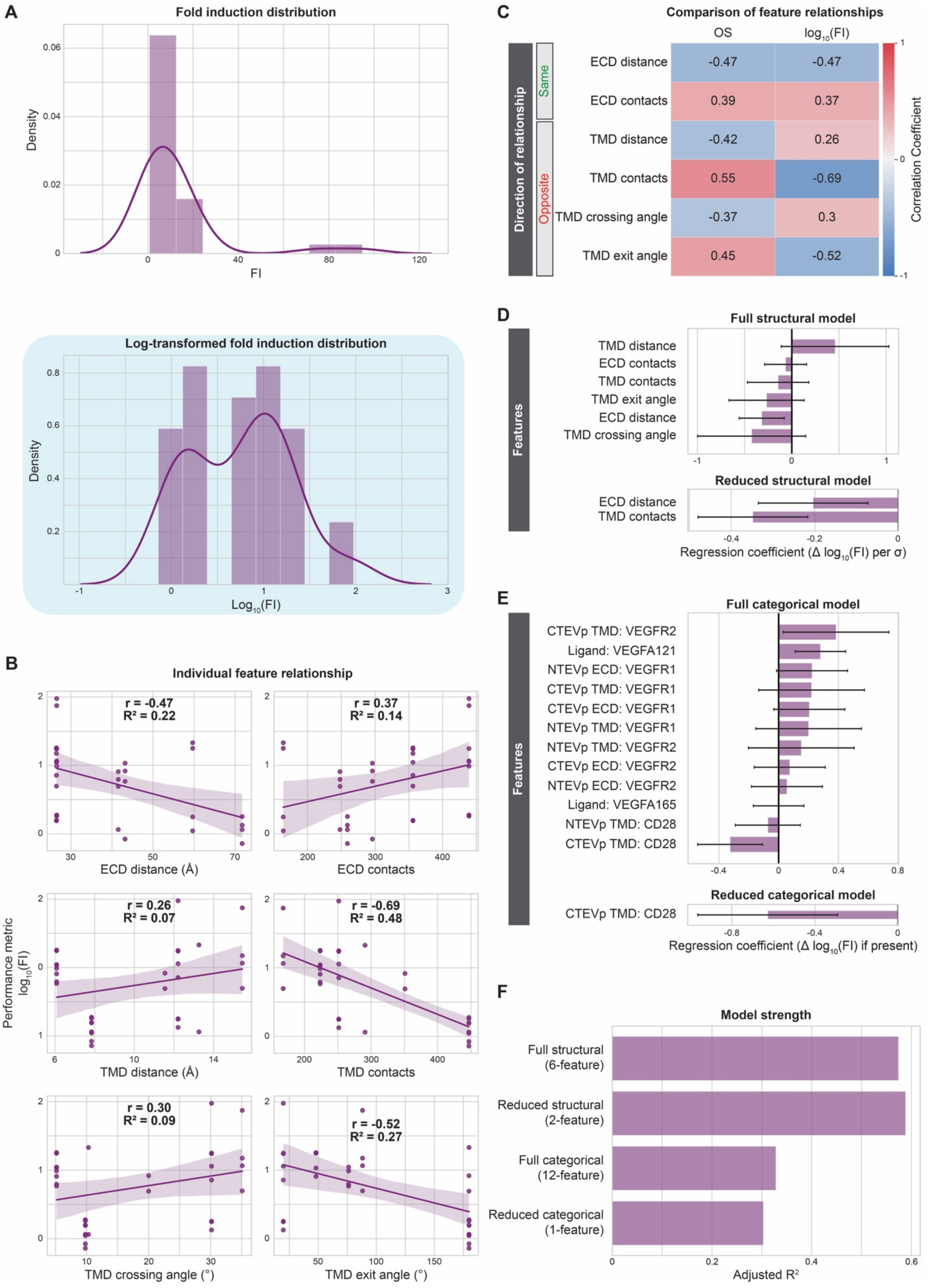

A
Fold induction distribution
Density
FI
Log-transformed fold induction distribution
Log10(FI)
B
Individual feature relationship
Performance metric log10(FI)
r = -0.47
R² = 0.22
ECD distance (Å)
r = 0.37
R² = 0.14
ECD contacts
r = 0.26
R² = 0.07
TMD distance (Å)
r = -0.69
R² = 0.48
TMD contacts
r = 0.30
R² = 0.09
TMD crossing angle (°)
r = -0.52
R² = 0.27
TMD exit angle (°)
C
Comparison of feature relationships
OS
log10(FI)
Direction of relationship
Same
Opposite
ECD distance -0.47 -0.47
ECD contacts 0.39 0.37
TMD distance -0.42 0.26
TMD contacts 0.55 -0.69
TMD crossing angle -0.37 0.3
TMD exit angle 0.45 -0.52
Correlation Coefficient
D
Full structural model
Features
TMD distance
ECD contacts
TMD contacts
TMD exit angle
ECD distance
TMD crossing angle
Reduced structural model
ECD distance
TMD contacts
Regression coefficient (Δ log10(FI) per σ)
E
Full categorical model
Features
CTEVp TMD: VEGFR2
Ligand: VEGFA121
NTEVp ECD: VEGFR1
CTEVp TMD: VEGFR1
CTEVp ECD: VEGFR1
NTEVp TMD: VEGFR1
NTEVp TMD: VEGFR2
CTEVp ECD: VEGFR2
NTEVp ECD: VEGFR2
Ligand: VEGFA165
NTEVp TMD: CD28
CTEVp TMD: CD28
Reduced categorical model
CTEVp TMD: CD28
Regression coefficient (Δ log10(FI) if present)
F
Model strength
Full structural (6-feature)
Reduced structural (2-feature)
Full categorical (12-feature)
Reduced categorical (1-feature)
Adjusted R²

**Supplementary Figure 2: Structural features account for considerable variation in VEGF MESA fold induction**

(**A**) The distribution of VEGF MESA FIs (top) and log-transformed FIs (bottom) with a fitted kernel density estimate (KDE). Given the right skew of the FI distribution, we selected log-transformed FI as the performance metric for further analysis. (**B**) Evaluation of the linear relationship between each individual structural feature and log-transformed FI. (**C**) Comparison between the sign of the relationship between each structural feature with OS and $\log_{10}$(FI) as the performance metric. Both ECD metrics correlate in the same direction for either performance metric, while all four TMD features correlate in opposite directions with the two performance metrics. (**D**) Magnitude of the coefficients for the full (six-feature) and reduced (two-feature) structural models. Error bars represent +/- 95% confidence intervals calculated using Student's t-distribution. (**E**) Magnitude of the coefficients for the full (twelve-feature) and reduced (two-feature) categorical models. Error bars represent +/- 95% confidence intervals calculated using Student's t-distribution. (**F**) Comparison of the four models' strengths in explaining variation in OS, as determined by adjusted $R^2$. Abbreviations: vascular endothelial growth factor receptor (VEGFR), vascular endothelial growth factor receptor 1 (VEGFR1), vascular endothelial growth factor receptor 2 (VEGFR2), vascular endothelial growth factor (VEGF), adjusted $R^2$ (Adj. $R^2$), Pearson's correlation coefficient (r), coefficient of determination ($R^2$), on-state reporter expression (OS), fold induction (FI), ectodomain (ECD), transmembrane domain (TMD), cluster of differentiation 28 (CD28), N-terminal half of Tobacco Etch Virus Protease (NTEVp), C-terminal half of Tobacco Etch Virus Protease (CTEVp), change in ($\Delta$), standard deviation ($\sigma$).

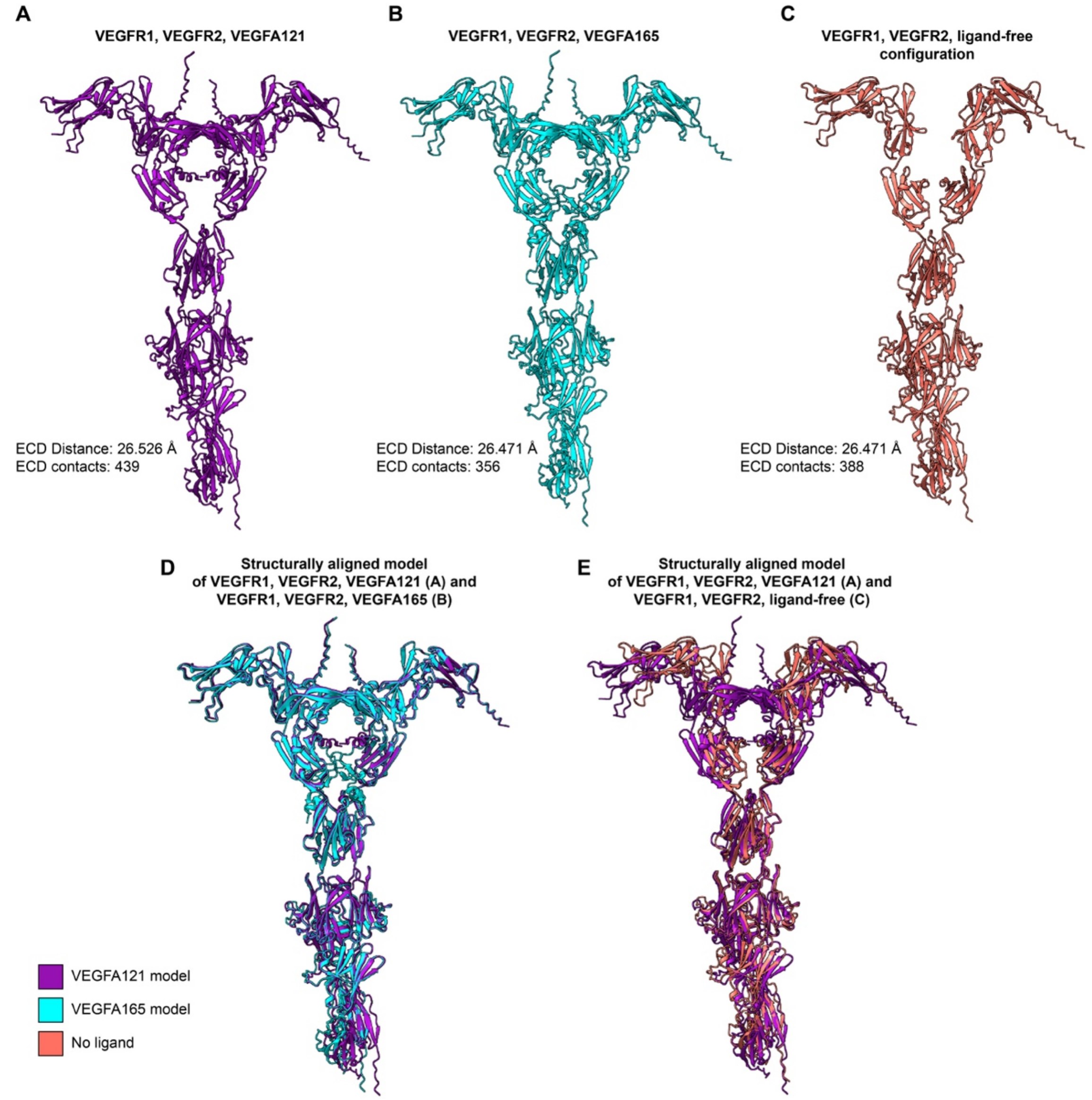


**Supplementary Figure 3: Comparison of ColabFold-generated VEGFR ECD models**

(**A**) Structural model of VEGFR1 and VEGFR2 in complex with a VEGFA121 dimer (**Supplementary Data 1, Structure 6**). (**B**) Structural model of VEGFR1 and VEGFR2 in complex with a VEGFA165 dimer (**Supplementary Data 1, Structure 7**). (**C**) Structural model of VEGFR1 and VEGFR2 dimerized in the absence of ligand (apo configuration) (**Supplementary Data 1, Structure 19**). (**D**) Superimposition of the structural model in **A** and **B**, wherein the difference in the structures is the ligand: VEGFA121 or VEGFA165. The structures have a high degree of similarity, resulting in similar measurements for their ECD features. (**E**) Superimposition of the structural model in **A** and **C**, wherein the difference in the structures is the ligand: VEGFA121 or apo (no ligand). The structures have a high degree of similarity, resulting in highly similar measurements for their ECD features. Abbreviations: vascular endothelial growth factor receptor 1 (VEGFR1), vascular endothelial growth factor receptor 2 (VEGFR2), vascular endothelial growth factor A, 121 amino acid isoform (VEGFA121), vascular endothelial growth factor A, 165 amino acid isoform (VEGFA165).

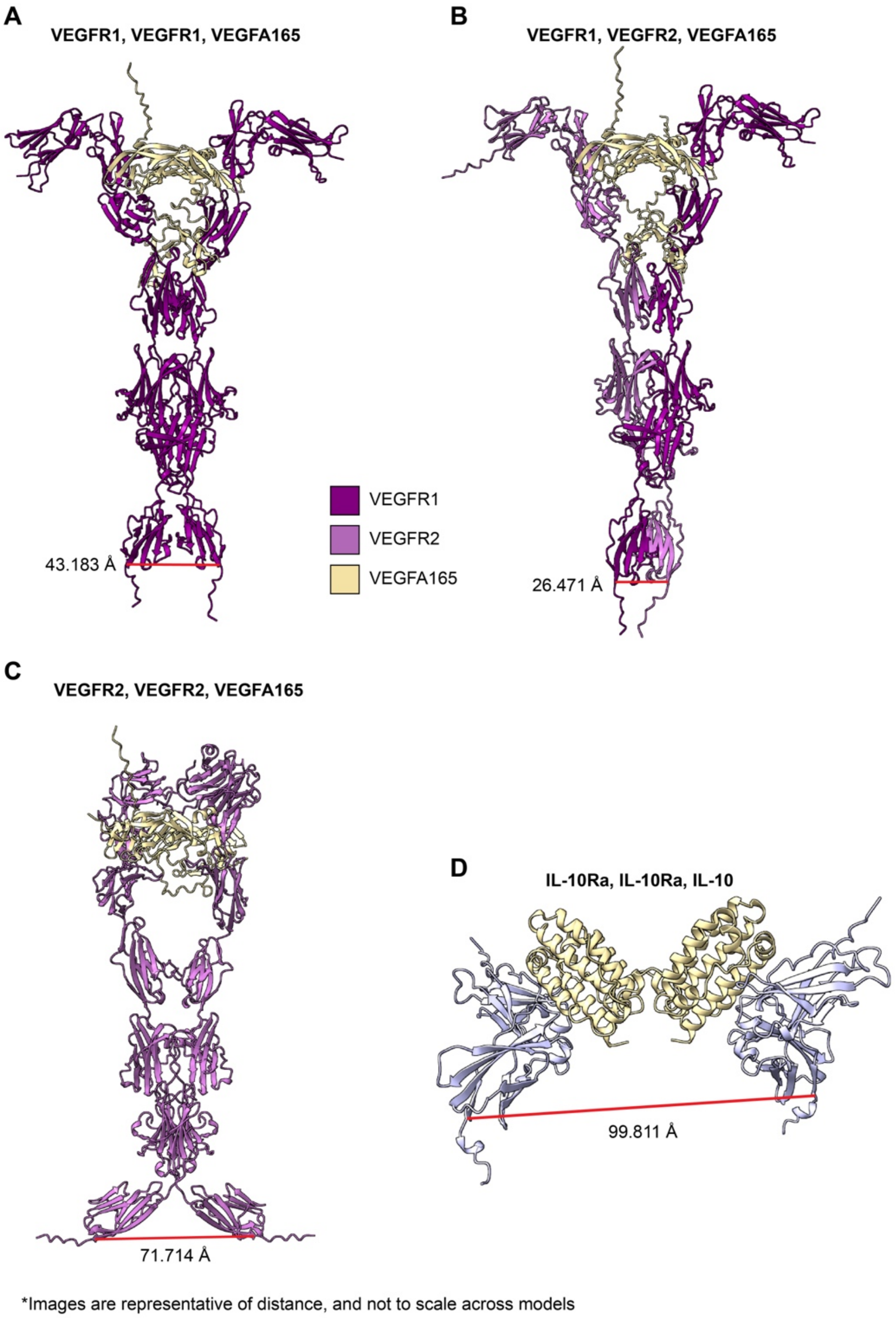


**Supplementary Figure 4: ECD distance varies considerably between ColabFold-generated models**
(**A**) ECD distance measurement of the structural model for VEGFR1 and VEGFR1 in complex with a VEGFA165 dimer (**Supplementary Data 1, Structure 5**). (**B**) ECD distance measurement of the structural model for VEGFR1 and VEGFR2 in complex with a VEGFA165 dimer (**Supplementary Data 1, Structure 7**). (**C**) ECD distance measurement of the structural model for VEGFR2 and VEGFR2 in complex with a VEGFA165 dimer (**Supplementary Data 1, Structure 9**). (**D**) ECD distance measurement of the structural model for IL-10Ra and IL-10Ra in complex with an IL-10 dimer (**Supplementary Data 1, Structure 13**). Abbreviations: ectodomain (ECD), vascular endothelial growth factor receptor 1 (VEGFR1), vascular endothelial growth factor receptor 2 (VEGFR2), vascular endothelial growth factor A, 121 amino acid variant (VEGFA121), vascular endothelial growth factor A, 165 amino acid variant (VEGFA165), interleukin 10 receptor a (IL-10Ra), interleukin 10 (IL-10).

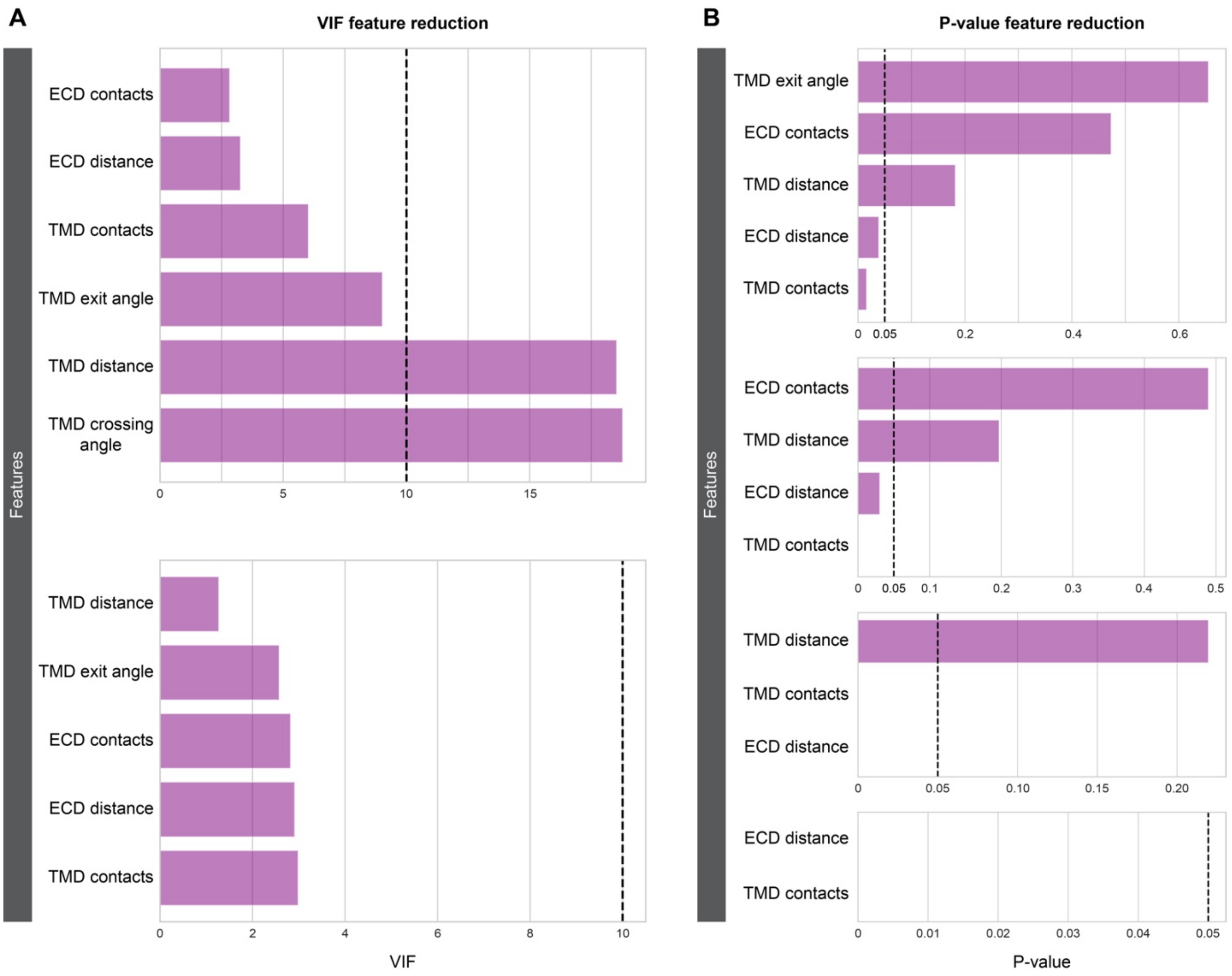


**Supplementary Figure 5**: **VEGFR structural model reduction methodology**
(**A**) Variance inflation factor (VIF) calculation for all six structural features (top). Removing TMD crossing angle resulted in lowering the VIF for all features below the threshold of 10. (**B**) Backwards elimination for the remaining five structural features (top), removing one feature each round based upon the coefficient p-value (α = 0.05), until only statistically significant features remain (bottom). Abbreviations: variance inflation factor (VIF), ectodomain (ECD), transmembrane domain (TMD).

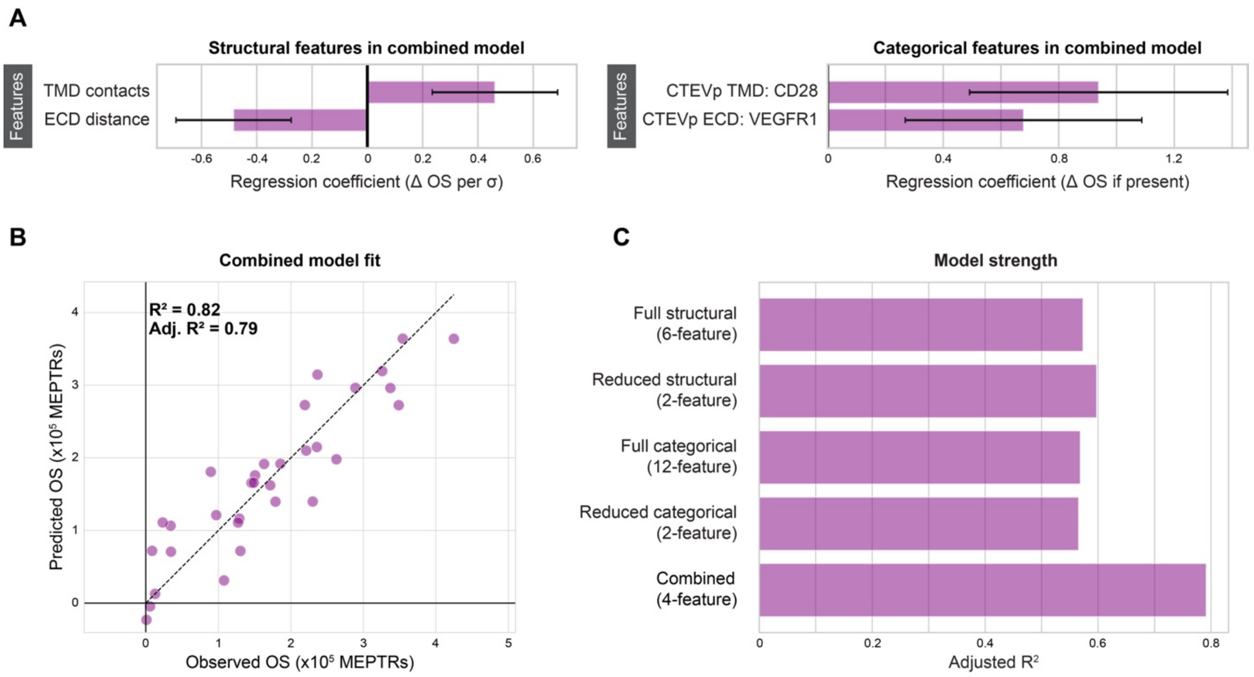


**Supplementary Figure 6: VEGF MESA combined model improves explanatory strength**

(**A**) Magnitude of the structural coefficients (left) and the categorical features (right) in the combined (four-feature) model. Error bars represent +/- 95% confidence intervals calculated using Student's t-distribution. (**B**) Combined (four-feature) model fit to explain variation in VEGF MESA OS. (**C**) Comparison of the four models' strengths in explaining variation in OS, as determined by adjusted $R^2$. Combining the features from the reduced structural model and reduced categorical model yielded a combined model with greater explanatory strength, indicating that the structural features and categorical features do not directly encode the same information to explain variation in OS. Abbreviations: transmembrane domain (TMD), ectodomain (ECD), C-terminal half of Tobacco Etch Virus Protease (CTEVp), cluster of differentiation 28 (CD28), adjusted $R^2$ (Adj. $R^2$), Pearson's correlation coefficient (r), coefficient of determination ($R^2$), on-state reporter expression (OS), molecules of equivalent PE-TexasRed (MEPTRs), change in (Δ), standard deviation (σ).

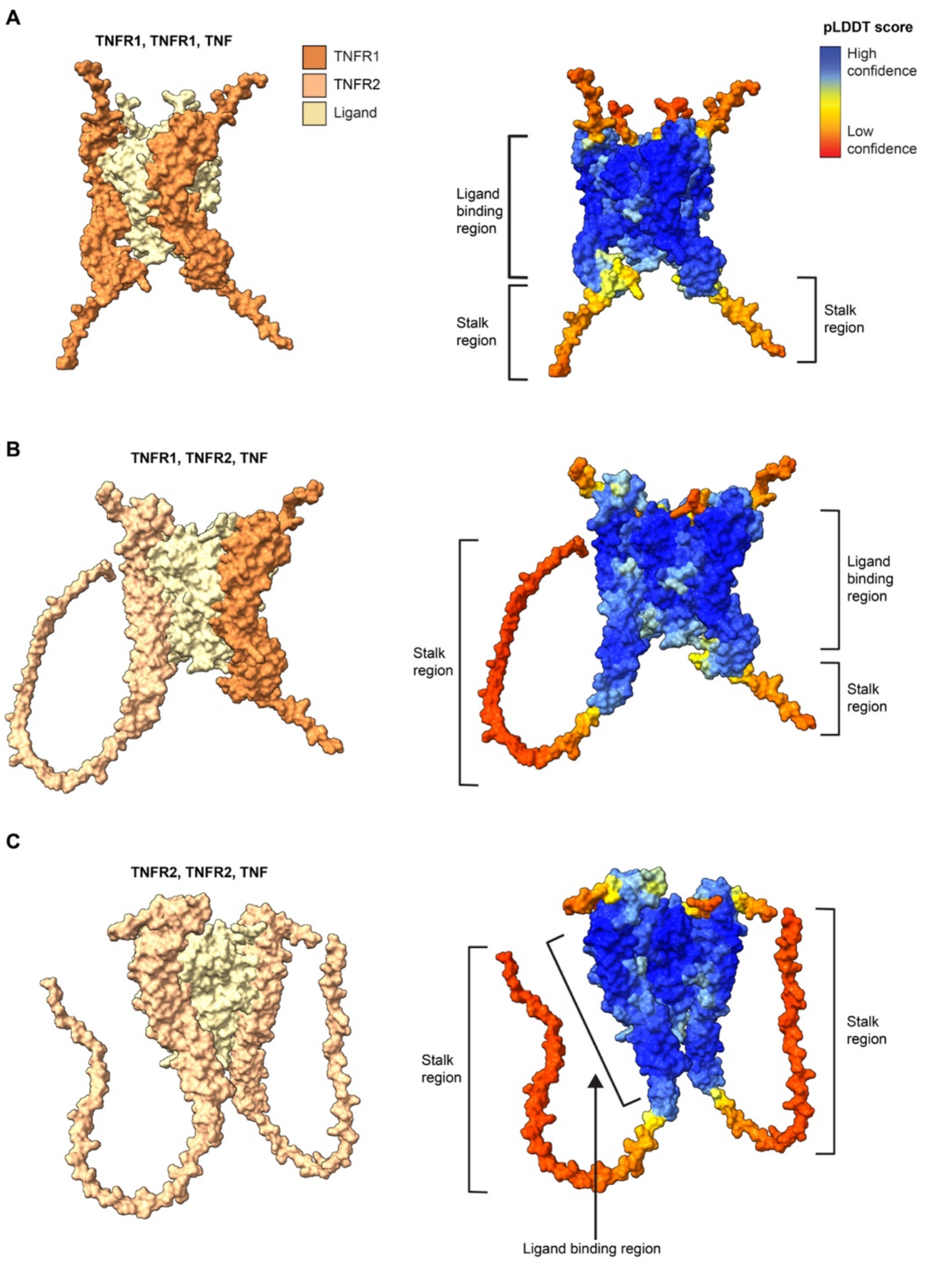


**Supplementary Figure 7: TNFR stalk regions are unstructured and have low confidence scores**
(**A**) Structural model of a TNFR1 and TNFR1 in complex with TNF colored by protein (left) and confidence score (right) (**Supplementary Data 1, Structure 10**). The stalk regions have low confidence scores. (**B**) Structural model of a TNFR1 and TNFR2 in complex with TNF colored by protein (left) and confidence score (right) (**Supplementary Data 1, Structure 11**). The stalk regions have low confidence scores and the TNFR2 stalk region folds upward, toward the ligand-binding region. (**C**) Structural model of a TNFR2 and TNFR2 in complex with TNF colored by protein (left) and confidence score (right) (**Supplementary Data 1, Structure 12**). The stalk regions have low confidence scores and both TNFR2 stalk region folds upward, toward the ligand-binding region. Abbreviations: predicted local distance difference test (pLDDT), tumor necrosis factor receptor (TNFR), tumor necrosis factor receptor 1 (TNFR1), tumor necrosis factor receptor 2 (TNFR2), tumor necrosis factor (TNF).

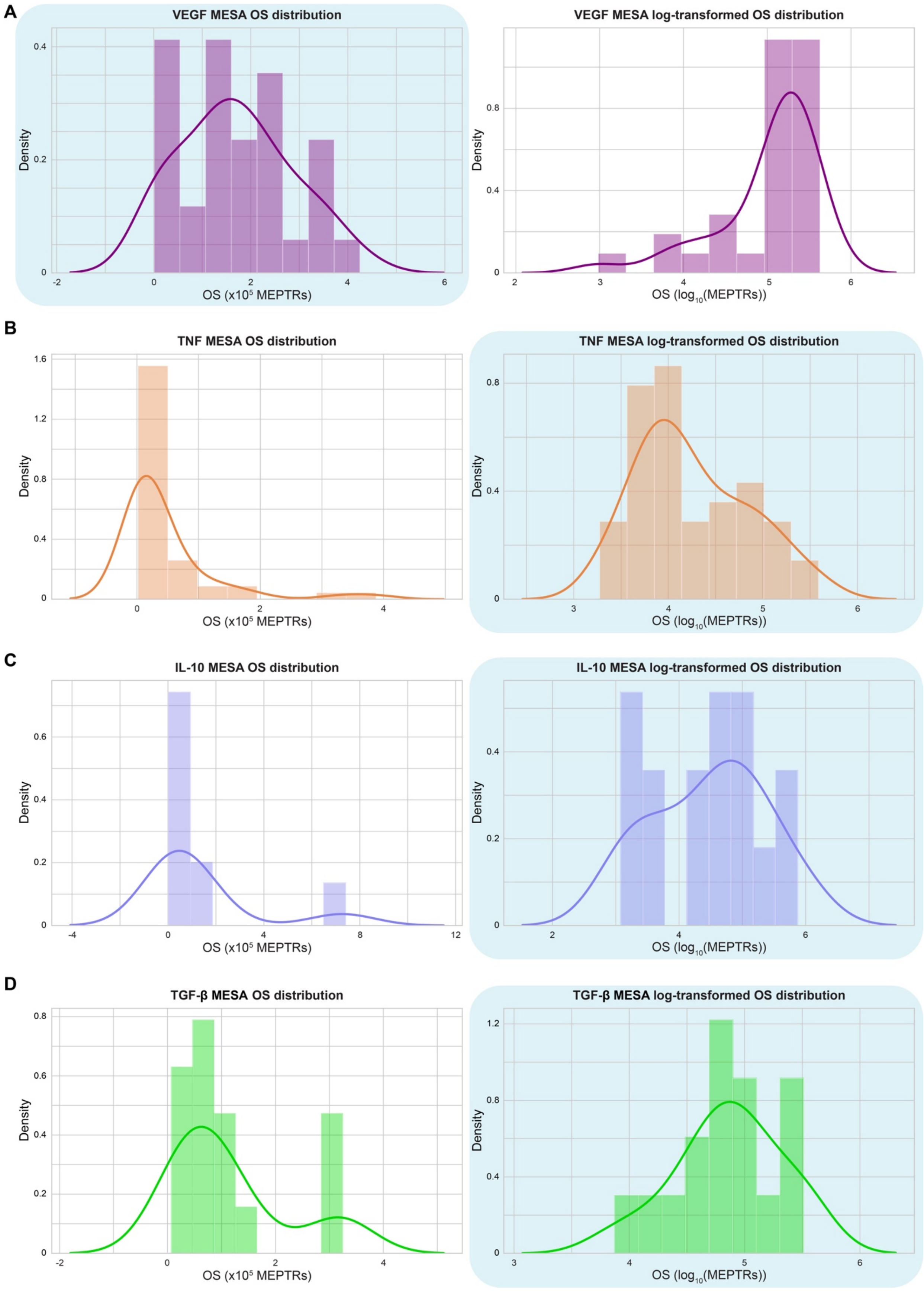

A
VEGF MESA OS distribution
Density
OS (x10^5 MEPTRs)
VEGF MESA log-transformed OS distribution
Density
OS (log10(MEPTRs))
B
TNF MESA OS distribution
Density
OS (x10^5 MEPTRs)
TNF MESA log-transformed OS distribution
Density
OS (log10(MEPTRs))
C
IL-10 MESA OS distribution
Density
OS (x10^5 MEPTRs)
IL-10 MESA log-transformed OS distribution
Density
OS (log10(MEPTRs))
D
TGF-β MESA OS distribution
Density
OS (x10^5 MEPTRs)
TGF-β MESA log-transformed OS distribution
Density
OS (log10(MEPTRs))

**Supplementary Figure 8**: **Selection of OS distributions**

(**A**) Distribution of VEGF MESA OS (left) and log-transformed OS (right) with fitted kernel density estimate (KDE). OS was selected as the performance metric for further analysis as log-transformed OS is left-skewed. (**B**) Distribution of TNF MESA OS (left) and log-transformed OS (right) with fitted KDEs. Log-transformed OS was selected as the performance metric for further analysis as OS is right-skewed. (**C**) Distribution of IL-10 MESA OS (left) and log-transformed OS (right) with fitted KDEs. Log-transformed OS was selected as the performance metric for further analysis as OS is right-skewed. (**D**) Distribution of TGFβ MESA OS (left) and log-transformed OS (right) with fitted KDEs. Log-transformed OS was selected as the performance metric for further analysis as OS is right-skewed. Abbreviations: on-state reporter expression (OS), vascular endothelial growth factor (VEGF), tumor necrosis factor (TNF), interleukin 10 (IL-10), transforming growth factor β (TGFβ), modular extracellular sensor architecture (MESA), ectodomain (ECD), transmembrane domain (TMD), molecules of equivalent PE-TexasRed (MEPTRs).

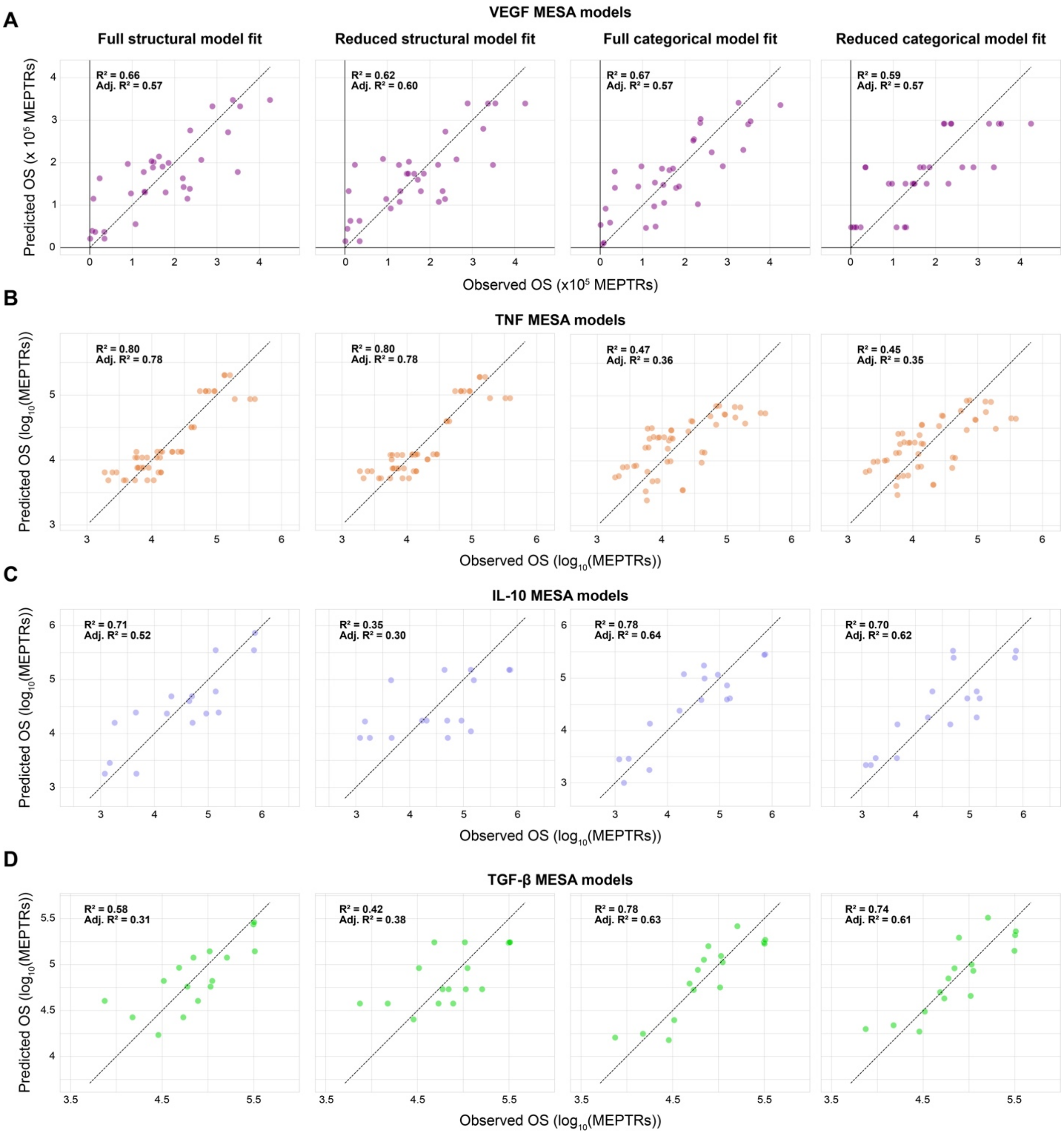


**Supplementary Figure 9: Comparison between structural and categorical model fit**

(**A**) Comparison between the VEGF MESA full structural model (six-feature) (left), reduced structural model (two-feature) (center left), full categorical model (twelve-feature) (center right), and the reduced categorical model (two-feature) (right) fit to explain variation in OS. (**B**) Comparison between the TNF MESA full structural model (six-feature) (left), reduced structural model (three-feature) (center left), full categorical model (thirteen-feature) (center right), and the reduced categorical model (eleven-feature) (right) fit to explain variation in log-transformed OS. (**C**) Comparison between the IL-10 MESA full structural model (six-feature) (left), reduced structural model (one-feature) (center left), full categorical model (ten-feature) (center right), and the reduced categorical model (five-feature) (right) fit to explain variation in log-transformed OS. (**D**) Comparison between the TGFβ MESA full structural model (six-feature) (left), reduced structural model (one-feature) (center left), full categorical model (ten-feature) (center right), and the reduced categorical model (eight-feature) (right) fit to explain variation in log-transformed OS. Abbreviations: on-state reporter expression (OS), vascular endothelial growth factor (VEGF), tumor necrosis factor (TNF), interleukin 10 (IL-10), transforming growth factor β (TGFβ), modular extracellular sensor architecture (MESA), ectodomain (ECD), transmembrane domain (TMD), Pearson's correlation coefficient (r), adjusted $R^2$ (Adj. $R^2$), coefficient of determination ($R^2$), molecules of equivalent PE-TexasRed (MEPTRs).

**Supplementary Table 1: ColabFold-generated structural models and confidence scores used in this study[1].**

| Model ID[2] | Receptor chains | Ligand | pLDDT score | pTM score | ipTM score |
|---|---|---|---|---|---|
| Structure 1 | VEGFR2 NTEVp, VEGFR1 CTEVp | VEGFA165 (dimer) | 72.2 | 0.601 | 0.522 |
| Structure 2 | IL-10Rb NTEVp, IL-10Ra CTEVp | IL-10 (dimer) | 85.8 | 0.702 | 0.688 |
| Structure 3 | IL-10Rb NTEVp (CD28 TMD), IL-10Ra CTEVp (CD28TMD) | IL-10 (dimer) | 80.5 | 0.656 | 0.652 |
| Structure 4 | VEGFR1, VEGFR1 | VEGFA121 (dimer) | 85.8 | 0.703 | 0.689 |
| Structure 5 | VEGFR1, VEGFR1 | VEGFA165 (dimer) | 85.2 | 0.682 | 0.678 |
| Structure 6 | VEGFR1, VEGFR2 | VEGFA121 (dimer) | 84.2 | 0.678 | 0.663 |
| Structure 7 | VEGFR1, VEGFR2 | VEGFA165 (dimer) | 82.2 | 0.665 | 0.658 |
| Structure 8 | VEGFR2, VEGFR2 | VEGFA121 (dimer) | 81.8 | 0.638 | 0.62 |
| Structure 9 | VEGFR2, VEGFR2 | VEGFA165 (dimer) | 80.1 | 0.636 | 0.628 |
| Structure 10 | TNFR1, TNFR1 | TNF (trimer) | 89.8 | 0.899 | 0.885 |
| Structure 11 | TNFR1, TNFR2 | TNF (trimer) | 87 | 0.87 | 0.851 |
| Structure 12 | TNFR2, TNFR2 | TNF (trimer) | 85.1 | 0.85 | 0.827 |
| Structure 13 | IL-10Ra, IL-10Ra | IL-10 (dimer) | 91.4 | 0.869 | 0.865 |
| Structure 14 | IL-10Ra, IL-10Ra, IL-10Rb, IL-10Rb | IL-10 (dimer) | 89.1 | 0.853 | 0.855 |
| Structure 15 | IL-10Rb, IL-10Rb | IL-10 (dimer) | 91.5 | 0.78 | 0.74 |
| Structure 16 | TGFβR1, TGFβR1 | TGF-β (dimer) | 80.9 | 0.737 | 0.676 |
| Structure 17 | TGFβR1, TGFβR1, TGFβR2, TGFβR2 | TGF-β (dimer) | 84.1 | 0.787 | 0.759 |
| Structure 18 | TGFβR2, TGFβR2 | TGF-β (dimer) | 81.4 | 0.633 | 0.571 |
| Structure 19 | VEGFR1, VEGFR2 | apo (no ligand) | 82.2 | 0.599 | 0.608 |

1. Abbreviations: Model identification (Model ID), ectodomains (ECDs), predicted local distance difference test (pLDDT), predicted template modeling (pTM), interfacial predicted template modeling (ipTM)
2. Individual model files are included in supporting materials online as **Supplementary Data 1**.

**Supplementary Table 2: Peptide sequences used in this study.**

| Sequence ID | Description: ECD, TMD (if present), ICD (if present) | Amino acid sequence |
|---|---|---|
| Seq 01 | VEGFR2 ECD, VEGFR2 TMD, NTEVp[1] | ASVGLPSVSLDLPRLSIQKDILTIKANTTLQITCRGQRDLDWLWPNN<br>QSGSEQRVEVTECSDGLFCKTLTIPKVIGNDTGAYKCFYRETDLAS<br>VIYVYVQDYRSPFIASVSDQHGVVYITENKNKTVVIPCLGSISNLNVS<br>LCARYPEKRFVPDGNRISWDSKKGFTIPSYMISYAGMVFCEAKIND<br>ESYQSIMYIVVVVGYRIYDVVLSPSHGIELSVGEKLVLNCTARTELNV<br>GIDFNWEYPSSKHQHKKLVNRDLKTQSGSEMKKFLSTLTIDGVTRS<br>DQGLYTCAASSGLMTKKNSTFVRVHEKPFVAFGSGMESLVEATVG<br>ERVRIPAKYLGYPPPEIKWYKNGIPLESNHTIKAGHVLTIMEVSERDT<br>GNYTVILTNPISKEKQSHVVSLVVYVPPQIGEKSLISPVDSYQYGTT<br>QTLTCTVYAIPPPHHIHWYWQLEEECANEPSQAVSVTNPYPCEEW<br>RSVEDFQGGNKIEVNKNQFALIEGKNKTVSTLVIQAANVSALYKCEA<br>VNKVGRGERVISFHVTRGPEITLQPDMQPTEQESVSLWCTADRST<br>FENLTWYKLGPQPLPIHVGELPTPVCKNLDTLWKLNATMFSNSTND<br>ILIMELKNASLQDQGDYVCLAQDRKTKKRHCVVRQLTVLERVAPTIT<br>GNLENQTTSIGESIEVSCTASGNPPPQIMWFKDNETLVEDSGIVLKD<br>GNRNLTIRRVRKEDEGLYTCQACSVLGCAKVEAFFIIEGAQEKTNLE<br>IIILVGTAVIAMFFWLLLVIILRTVKGAESLFKGPRDYNPISSTICHLTN<br>ESDGHTTSLYGIGFGPFIITNKHLFRRNNGTLLVQSLHGVFKVKNTT<br>TLQQSLIDGRDMIIIRMPKDFPPFPQKLKFREPQREERICLVTTNFQT |
| Seq 02 | VEGFR1 ECD, VEGFR1 TMD, CTEVp[2] | SKLKDPELSLKGTQHIMQAGQTLHLQCRGEAAHKWSLPEMVSKES<br>ERLSITKSACGRNGKQFCSTLTLNTAQANHTGFYSCKYLAVPTSKK<br>KETESAIYIFISDTGRPFVEMYSEIPEIIHMTEGRELVIPCRVTSPNITV<br>TLKKFPLDTLIPDGKRIIWDSRKGFIISNATYKEIGLLTCEATVNGHLY<br>KTNYLTHRQTNTIIDVQISTPRPVKLLRGHTLVLNCTATTPLNTRVQ<br>MTWSYPDEKNKRASVRRRIDQSNSHANIFYSVLTIDKMQNKDKGLY<br>TCRVRSGPSFKSVNTSVHIYDKAFITVKHRKQQVLETVAGKRSYRL<br>SMKVKAFPSPEVVWLKDGLPATEKSARYLTRGYSLIIKDVTEEDAG<br>NYTILLSIKQSNVFKNLTATLIVNVKPQIYEKAVSSFPDPALYPLGSR<br>QILTCTAYGIPQPTIKWFWHPCNHNHSEARCDFCSNNEESFILDAD<br>SNMGNRIESITQRMAIIEGKNKMASTLVVADSRISGIYICIASNKVGTV<br>GRNISFYITDVPNGFHVNLEKMPTEGEDLKLSCTVNKFLYRDVTWIL<br>LRTVNNRTMHYSISKQKMAITKEHSITLNLTIMNVSLQDSGTYACRA<br>RNVYTGEEILQKKEITIRDQEAPYLLRNLSDHTVAISSSTTLDCHANG<br>VPEPQITWFKNNHKIQQEPGIILGPGSSTLFIERVTEEDEGVYHCKA<br>TNQKGSVESSAYLTVQGTSDKSNLELITLTCTCVAATLFWLLLTLFIR<br>KMKRGAKSMSSMVSDTSCTFPSSDGIFWKHWIQTKDGQCGSPLV<br>STRDGFIVGIHSASNFTNTNNYFTSVPKNFMELKTNQEAQQWVSG<br>WRLNADSVLWGGHKVFMVKPEEPFQPVKEATQLMN |
| Seq 03 | VEGFA121 (monomer) | APMAEGGGQNHHEVVKFMDVYQRSYCHPIETLVDIFQEYPDEIEYI<br>FKPSCVPLMRCGGCCNDEGLECVPTEESNITMQIMRIKPHQGQHIG<br>EMSFLQHNKCECRPKKDRARQEKCDKPRR |
| Seq 04 | VEGFA165 (monomer) | APMAEGGGQNHHEVVKFMDVYQRSYCHPIETLVDIFQEYPDEIEYI<br>FKPSCVPLMRCGGCCNDEGLECVPTEESNITMQIMRIKPHQGQHIG<br>EMSFLQHNKCECRPKKDRARQENPCGPCSERRKHLFVQDPQTCK<br>CSCKNTDSRCKARQLELNERTCRCDKPRR |

| | | |
|---|---|---|
| Seq 05 | IL-10Rb ECD, IL-10Rb TMD, NTEVp[1] | MVPPPENVRMNSVNFKNILQWESPAFAKGNLTFTAQYLSYRIFQDK<br>CMNTTLTECDFSSLSKYGDHTLRVRAEFADEHSDWVNITFCPVDDT<br>IIGPPGMQVEVLADSLHMRFLAPKIENEYETWTMKNVYNSWTYNVQ<br>YWKNGTDEKFQITPQYDFEVLRNLEPWTTYCVQVRGFLPDRNKAG<br>EWSEPVCEQTTHDETVPSWMVAVILMASVFMVCLALLGCFALLWC<br>VYKKGAESLFKGPRDYNPISSTICHLTNESDGHTTSLYGIGFGPFIIT<br>NKHLFRRNNGTLLVQSLHGVFKVKNTTTLQQSLIDGRDMIIIRMPKD<br>FPPFPQKLKFREPQREERICLVTTNFQT |
| Seq 06 | IL-10Ra ECD, IL-10Ra TMD, CTEVp[2] | HGTELPSPPSVWFEAEFFHHILHWTPIPNQSESTCYEVALLRYGIES<br>WNSISNCSQTLSYDLTAVTLDLYHSNGYRARVRAVDGSRHSNWTV<br>TNTRFSVDEVTLTVGSVNLEIHNGFILGKIQLPRPKMAPANDTYESIF<br>SHFREYEIAIRKVPGNFTFTHKKVKHENFSLLTSGEVGEFCVQVKPS<br>VASRSNKGMWSKEECISLTRQYFTVTNVIIFFAFVLLLSGALAYCLAL<br>QLYVRRRKKGAKSMSSMVSDTSCTFPSSDGIFWKHWIQTKDGQC<br>GSPLVSTRDGFIVGIHSASNFTNTNNYFTSVPKNFMELKTNQEAQQ<br>WVSGWRLNADSVLWGGHKVFMVKPEEPFQPVKEATQLMN |
| Seq 07 | IL-10 (monomer) | SPGQGTQSENSCTHFPGNLPNMLRDLRDAFSRVKTFFQMKDQLD<br>NLLLKESLLEDFKGYLGCQALSEMIQFYLEEVMPQAENQDPDIKAH<br>VNSLGENLKTLRLRLRRCHRFLPCENKSKAVEQVKNAFNKLQEKGI<br>YKAMSEFDIFINYIEAYMTMKIRN |
| Seq 08 | IL-10Rb ECD, CD28 TMD, NTEVp[1] | MVPPPENVRMNSVNFKNILQWESPAFAKGNLTFTAQYLSYRIFQDK<br>CMNTTLTECDFSSLSKYGDHTLRVRAEFADEHSDWVNITFCPVDDT<br>IIGPPGMQVEVLADSLHMRFLAPKIENEYETWTMKNVYNSWTYNVQ<br>YWKNGTDEKFQITPQYDFEVLRNLEPWTTYCVQVRGFLPDRNKAG<br>EWSEPVCEQTTHDETVPSTGLVVVAGVLFCYGLLVTVALCVGGGS<br>GGGAESLFKGPRDYNPISSTICHLTNESDGHTTSLYGIGFGPFIITNK<br>HLFRRNNGTLLVQSLHGVFKVKNTTTLQQSLIDGRDMIIIRMPKDFP<br>PFPQKLKFREPQREERICLVTTNFQT |
| Seq 09 | IL-10Ra ECD, CD28 TMD, CTEVp[2] | HGTELPSPPSVWFEAEFFHHILHWTPIPNQSESTCYEVALLRYGIES<br>WNSISNCSQTLSYDLTAVTLDLYHSNGYRARVRAVDGSRHSNWTV<br>TNTRFSVDEVTLTVGSVNLEIHNGFILGKIQLPRPKMAPANDTYESIF<br>SHFREYEIAIRKVPGNFTFTHKKVKHENFSLLTSGEVGEFCVQVKPS<br>VASRSNKGMWSKEECISLTRQYFTVTNTGLVVVAGVLFCYGLLVTV<br>ALCVGGGSGGGAKSMSSMVSDTSCTFPSSDGIFWKHWIQTKDGQ<br>CGSPLVSTRDGFIVGIHSASNFTNTNNYFTSVPKNFMELKTNQEAQ<br>QWVSGWRLNADSVLWGGHKVFMVKPEEPFQPVKEATQLMN |
| Seq 10 | VEGFR1 ECD | SKLKDPELSLKGTQHIMQAGQTLHLQCRGEAAHKWSLPEMVSKES<br>ERLSITKSACGRNGKQFCSTLTLNTAQANHTGFYSCKYLAVPTSKK<br>KETESAIYIFISDTGRPFVEMYSEIPEIIHMTEGRELVIPCRVTSPNITV<br>TLKKFPLDTLIPDGKRIIWDSRKGFIISNATYKEIGLLTCEATVNGHLY<br>KTNYLTHRQTNTIIDVQISTPRPVKLLRGHTLVLNCTATTPLNTRVQ<br>MTWSYPDEKNKRASVRRRIDQSNSHANIFYSVLTIDKMQNKDKGLY<br>TCRVRSGPSFKSVNTSVHIYDKAFITVKHRKQQVLETVAGKRSYRL<br>SMKVKAFPSPEVVWLKDGLPATEKSARYLTRGYSLIIKDVTEEDAG<br>NYTILLSIKQSNVFKNLTATLIVNVKPQIYEKAVSSFPDPALYPLGSR<br>QILTCTAYGIPQPTIKWFWHPCNHNHSEARCDFCSNNEESFILDAD<br>SNMGNRIESITQRMAIIEGKNKMASTLVVADSRISGIYICIASNKVGTV<br>GRNISFYITDVPNGFHVNLEKMPTEGEDLKLSCTVNKFLYRDVTWIL<br>LRTVNNRTMHYSISKQKMAITKEHSITLNLTIMNVSLQDSGTYACRA<br>RNVYTGEEILQKKEITIRDQEAPYLLRNLSDHTVAISSSTTLDCHANG<br>VPEPQITWFKNNHKIQQEPGIILGPGSSTLFIERVTEEDEGVYHCKA<br>TNQKGSVESSAYLTVQGTSDKSNLE |

| | | |
|---|---|---|
| Seq 11 | VEGFR2 ECD | ASVGLPSVSLDLPRLSIQKDILTIKANTTLQITCRGQRDLDWLWPNNQSGSEQRVEVTECSDGLFCKTLTIPKVIGNDTGAYKCFYRETDLASVIYVYVQDYRSPFIASVSDQHGVVYITENKNKTVVIPCLGSISNLNVSLCARYPEKRFVPDGNRISWDSKKGFTIPSYMISYAGMVFCEAKINDESYQSIMYIVVVVGYRIYDVVLSPSHGIELSVGEKLVLNCTARTELNVGIDFNWEYPSSKHQHKKLVNRDLKTQSGSEMKKFLSTLTIDGVTRSDQGLYTCAASSGLMTKKNSTFVRVHEKPFVAFGSGMESLVEATVGERVRIPAKYLGYPPPEIKWYKNGIPLESNHTIKAGHVLTIMEVSERDTGNYTVILTNPISKEKQSHVVSLVVYVPPQIGEKSLISPVDSYQYGTTQTLTCTVYAIPPPHHIHWYWQLEEECANEPSQAVSVTNPYPCEEWRSVEDFQGGNKIEVNKNQFALIEGKNKTVSTLVIQAANVSALYKCEAVNKVGRGERVISFHVTRGPEITLQPDMQPTEQESVSLWCTADRSTFENLTWYKLGPQPLPIHVGELPTPVCKNLDTLWKLNATMFSNSTNDILIMELKNASLQDQGDYVCLAQDRKTKKRHCVVRQLTVLERVAPTITGNLENQTTSIGESIEVSCTASGNPPPQIMWFKDNETLVEDSGIVLKDGNRNLTIRRVRKEDEGLYTCQACSVLGCAKVEAFFIIEGAQEKTNLE |
| Seq 12 | TNFR1 ECD | LVPHLGDREKRDSVCPQGKYIHPQNNSICCTKCHKGTYLYNDCPGPGQDTDCRECESGSFTASENHLRHCLSCSKCRKEMGQVEISSCTVDRDTVCGCRKNQYRHYWSENLFQCFNCSLCLNGTVHLSCQEKQNTVCTCHAGFFLRENECVSCSNCKKSLECTKLCLPQIENVKGTEDSGTT |
| Seq 13 | TNFR2 ECD | LPAQVAFTPYAPEPGSTCRLREYYDQTAQMCCSKCSPGQHAKVFCTKTSDTVCDSCEDSTYTQLWNWVPECLSCGSRCSSDQVETQACTREQNRICTCRPGWYCALSKQEGCRLCAPLRKCRPGFGVARPGTETSDVVCKPCAPGTFSNTTSSTDICRPHQICNVVAIPGNASMDAVCTSTSPTRSMAPGAVHLPQPVSTRSQHTQPTPEPSTAPSTSFLLPMGPSPPAEGSTGD |
| Seq 14 | TNF (monomer) | VRSSSRTPSDKPVAHVVANPQAEGQLQWLNRRANALLANGVELRDNQLVVPSEGLYLIYSQVLFKGQGCPSTHVLLTHTISRIAVSYQTKVNLLSAIKSPCQRETPEGAEAKPWYEPIYLGGVFQLEKGDRLSAEINRPDYLDFAESGQVYFGIIAL |
| Seq 15 | IL-10Ra ECD | HGTELPSPPSVWFEAEFFHHILHWTPIPNQSESTCYEVALLRYGIESWNSISNCSQTLSYDLTAVTLDLYHSNGYRARVRAVDGSRHSNWTVTNTRFSVDEVTLTVGSVNLEIHNGFILGKIQLPRPKMAPANDTYESIFSHFREYEIAIRKVPGNFTFTHKKVKHENFSLLTSGEVGEFCVQVKPSVASRSNKGMWSKEECISLTRQYFTVTN |
| Seq 16 | IL-10Rb ECD | MVPPPENVRMNSVNFKNILQWESPAFAKGNLTFTAQYLSYRIFQDKCMNTTLTECDFSSLSKYGDHTLRVRAEFADEHSDWVNITFCPVDDTIIGPPGMQVEVLADSLHMRFLAPKIENEYETWTMKNVYNSWTYNVQYWKNGTDEKFQITPQYDFEVLRNLEPWTTYCVQVRGFLPDRNKAGEWSEPVCEQTTHDETVPS |
| Seq 17 | TGFβR1 ECD | LQCFCHLCTKDNFTCVTDGLCFVSVTETTDKVIHNSMCIAEIDLIPRDRPFVCAPSSKTGSVTTTYCCNQDHCNKIELPTTVKSSPGLGPVEL |
| Seq 18 | TGFβR2 ECD | TIPPHVQKSVNNDMIVTDNNGAVKFPQLCKFCDVRFSTCDNQKSCMSNCSITSICEKPQEVCVAVWRKNDENITLETVCHDPKLPYHDFILEDAASPKCIMKEKKKPGETFFMCSCSSDECNDNIIFSEEYNTSNPDLLLVIFQ |
| Seq 19 | TGFβ1 (monomer) | ALDTNYCFSSTEKNCCVRQLYIDFRKDLGWKWIHEPKGYHANFCLGPCPYIWSLDTQYSKVLALYNQHNPGASAAPCCVPQALEPLPIVYYVGRKPKVEQLSNMIVRSCKCS |
| Seq 20 | VEGFR1 TMD | LITLTCTCVAATLFWLLLTLFI |
| Seq 21 | VEGFR2 TMD | IIILVGTAVIAMFFWLLLVII |
| Seq 22 | TNFR1 TMD | VLLPLVIFFGLCLLSLLFIGL |
| Seq 23 | TNFR2 TMD | FALPVGLIVGVTALGLLIIGVVNCVIMTQV |

| | | |
|---|---|---|
| Seq 24 | IL-10Ra TMD | VIIFFAFVLLLSGALAYCLAL |
| Seq 25 | IL-10Rb TMD | WMVAVILMASVFMVCLALLGCF |
| Seq 26 | TGFBR1 TMD | AAVIAGPVCFVCISLMLMVYI |
| Seq 27 | TGFBR2 TMD | VTGISLLPPLGVAISVIIIFY |
| Seq 28 | Murine CD28 TMD | LVVVAGVLFCYGLLVTVALCV |

1. The NTEVp variant used has an H75S mutation from wildtype
2. The CTEVp variant used has an L190K mutation from wildtype

**Supplementary Table 3: PREDDIMER TMD dimer quality scores**

| TMD 1 | TMD 2 | Fscor |
|---|---|---|
| VEGFR1 | VEGFR1 | 2.838 |
| VEGFR1 | VEGFR2 | 2.096 |
| VEGFR2 | VEGFR1 | 2.449 |
| VEGFR2 | VEGFR2 | 2.351 |
| VEGFR1 | CD28 | 2.645 |
| CD28 | VEGFR1 | 2.648 |
| VEGFR2 | CD28 | 2.448 |
| CD28 | VEGFR2 | 2.479 |
| TNFR1 | TNFR1 | 3.267 |
| TNFR1 | TNFR2 | 3.174 |
| TNFR2 | TNFR1 | 2.968 |
| TNFR2 | TNFR2 | 2.805 |
| TNFR1 | CD28 | 2.785 |
| CD28 | TNFR1 | 2.93 |
| TNFR2 | CD28 | 2.864 |
| CD28 | TNFR2 | 2.636 |
| IL-10Ra | IL-10Ra | 2.69 |
| IL-10Ra | IL-10Rb | 2.713 |
| IL-10Rb | IL-10Ra | 2.668 |
| IL-10Rb | IL-10Rb | 2.779 |
| IL-10Ra | CD28 | 2.719 |
| CD28 | IL-10Ra | 2.831 |
| IL-10Rb | CD28 | 2.317 |
| CD28 | IL-10Rb | 2.436 |
| TGFBR1 | TGFBR1 | 2.447 |
| TGFBR1 | TGFBR2 | 3.105 |
| TGFBR2 | TGFBR1 | 3.345 |
| TGFBR2 | TGFBR2 | 4.334 |
| TGFBR1 | CD28 | 2.815 |
| CD28 | TGFBR1 | 2.759 |
| TGFBR2 | CD28 | 3.549 |
| CD28 | TGFBR2 | 3.618 |
| CD28 | CD28 | 2.723 |
